# CeDNe: A multi-scale computational framework for modeling structure-function relationships in the *C. elegans* nervous system

**DOI:** 10.1101/2025.11.03.683805

**Authors:** Sahil Moza, Yun Zhang

**Affiliations:** Department of Organismic and Evolutionary Biology, Harvard University, Cambridge, MA, USA

## Abstract

Understanding how neural circuits generate behavior requires integrating structural and functional data across scales. *C. elegans* with its complete connectome, genetically identifiable neurons, single-cell transcriptome, neuropeptide-receptor distribution, and an amenability to simultaneous measurement of brain-wide neural activity and behavior presents a unique opportunity for such a multiscale circuit analysis. However, the absence of a unifying framework to connect these diverse datasets limits our ability to connect network structure and attributes with function. Here we introduce CeDNe (*C. elegans* Dynamical Network), an open-source computational framework that integrates anatomical, molecular, and imaging datasets into a unified graph-based representation that enables multimodal data analysis by cross-referencing different omics layers in a single computational environment. Specifically, CeDNe provides modular tools for visualizing and analyzing network connectivity, motif distribution, and circuit paths. Further, it incorporates a computational framework that simulates neural dynamics and optimizes network models to bridge structural connectivity with neural activity. Thus, CeDNe establishes a scalable foundation for data-driven modeling of the nervous system. This open-source tool not only facilitates computational connectomics and multimodal analyses in *C. elegans* but also serves as a generalizable framework for investigating structure-function relationships in neural networks of other organisms.

## Introduction

One of the central goals of neuroscience is to understand how the connectivity and properties of a nervous system shape information processing to direct behavior. In his Nobel Lecture “Nature’s gift to science”, Sydney Brenner envisioned the next frontier of neuroscience as CellMaps - detailed multidimensional computational maps that systematically integrate molecular, anatomical and functional information [1] to understand nervous system organization and operation. Since then, numerous organism-level atlases have emerged for *C. elegans*, spanning singlecell gene expression [2, 3], intercellular signal profiling (neurotransmitter [4, 5] and neuropeptide-receptor interactions [6]), and functional calcium imaging [7–12]. These mega-datasets in combination with the fully mapped wiring diagram [13–15], present a timely opportunity to develop a holistic understanding of the function of the nervous system [16]. However, the lack of a scalable framework that enables integrative analysis of anatomical, molecular, and functional datasets and connects these network attributes with dynamic modeling, limits our ability to achieve this goal. To address this unmet demand, we have developed **CeDNe** [s’Idni] ***C.elegans*** **D**ynamical **Ne**twork, a Python based platform that unifies diverse datasets, organizes them into a graph structure and constructs an evolving top-down computational model of the *C. elegans* nervous system at the multiscale level- from organism scale to molecular scale. CeDNe bridges the gap between genes, wiring diagram and neural function by incorporating the mega-datasets [13–15], [4, 5], [6],[2, 3], [7] and computational simulations into a graph-based architecture that enables network visualization and analysis, motif discovery, and predictive modeling. The unique strengths of CeDNe are that it enables simultaneous access and analysis of multiple types of omics and functional imaging data, integrates them with computational tools to generate testable predictions of network function and properties, and thus reduces the gap between neural data analysis and modeling.

Within the framework of CeDNe, the anatomical connectome is represented as a graph-based backbone with each node, connection and the whole graph serving as integrating points for multimodal sources of information. This approach effectively embeds the various types of data into a common multi-relational graph, that can both be used as background to store new information and also facilitates analyzing functional computations and dynamic model building. This enables researchers to test hypotheses on both existing and newly acquired data.

Here we use CeDNe to address fundamental questions about the core organizational principles of the *C. elegans* nervous system. First, how does network structure shape the flow of information? By exploring motifs in the *C. elegans* network, we find that chains of feed-forward loop motifs [17] are overrepresented and strongly aligned with the sensory-to-motor direction of information flow. Next, how is the expression of neurotransmitters distributed in the neural network subserving information flow from sensory to motor neurons? Using connectivity, ligand and receptor expression, we predict neurotransmitter identity for chemical synapses. We find that neurotransmitters segregate by identity in feed-forward loop chains from glutamatergic in the early layers to cholinergic in the deeper layers. Then, we ask how information propagates over the length of the sequential hierarchy. We simulate sequential hierarchies with a range of parameters and show that these simple chains can be useful for temporal patterning of responses to inputs. We further integrate the neuropeptidergic connectome to predict enriched neuropeptides in pharyngeal neurons, which can present specific neuropeptide-mediated volume transmissions as potential candidates for experimental investigation. Finally, we demonstrate the applicability of CeDNe to analysis of other systems, such as the nervous systems of male *C. elegans* and adult *Drosophila melanogaster*.

Thus, by simultaneously accessing and analyzing different datasets on various aspects of the nervous system and connecting with computational simulations, CeDNe leverages the ability to discover mechanistic relationships among independently acquired network attributes to uncover new insights that an individual resource alone cannot find. We propose that CeDNe complements the excellent platforms for the *bottom-up* biophysical modeling of the *C. elegans* nervous system [18, 19] and enables an opportunity to build towards coherent theories of brain function, which requires integration, collective interpretation, and continuous building upon existing knowledge.

## Results

### CeDNe: A unified framework for multi-scale circuit analysis

CeDNe organizes the *C. elegans* nervous system as a multi-layered directed graph, where nodes correspond to *Neurons* and edges encode different types of neuronal *Connections* including chemical synapses, gap-junctions, neuropeptidergic or functional interactions inferred from neural activity data (**Fig.1a**). Each component of the network can hold a range of biological properties. *Neurons* store data on cellular properties, such as their cell identity and gene expression (including neurotransmitters, neuropeptides, and receptors), and neural activities; *Connections* store information on connection properties, such as synaptic weights, connected neurons and neurochemistry inferred from single-cell transcriptomes **(Fig.1b**). Neurons can be grouped into one or more *NeuronGroup*s and Connections into one or more *ConnectionGroup*s, which can both hold relevant properties themselves. For example, all neurons that express a particular gene and all dopaminergic connections can belong to *Neuron-Group* and *ConnectionGroup* classes respectively. A *Path*, a series of non-repeating edges of a specific length connecting one neuron to another across the network, can be derived. New properties can be added to any of the components, allowing contextualization of new data with available data. This integrated representation builds the structural, neurochemical, and functional properties into a cohesive framework to support multi-modal analyses and modeling. Users can examine specific motifs, extract sub-networks of interest, and apply network theory approaches to identify functional features, including highly connected hubs, information flow pathways, and motif-enriched regions (**Fig.1c**). Subgraph extraction for motif analysis can allow users to isolate parts of their graph to analyze structural properties or integrate their own data in isolated *NeuronGroup*s or *ConnectionGroup*s of interest (**Fig.1c**). Next, CeDNe facilitates users to create functional top-down models from data and simulate their circuits of interest (**Fig.1d**). To facilitate users’ queries, CeDNe also provides multiple visualization and data exploration tools. Grouped and anatomical layouts can allow users to visualize rich information at the level of the whole nervous system(**Fig.1e,f** ). Other layouts such as shell layout can be used to look at sub-circuit connectivity and paths between neurons (**Fig.1g**). While currently optimized for *C. elegans*, its modular structure allows adaptation to other small nervous systems, such as *Drosophila* as demonstrated in later sections and regions of mammalian brains.

**Fig. 1.**
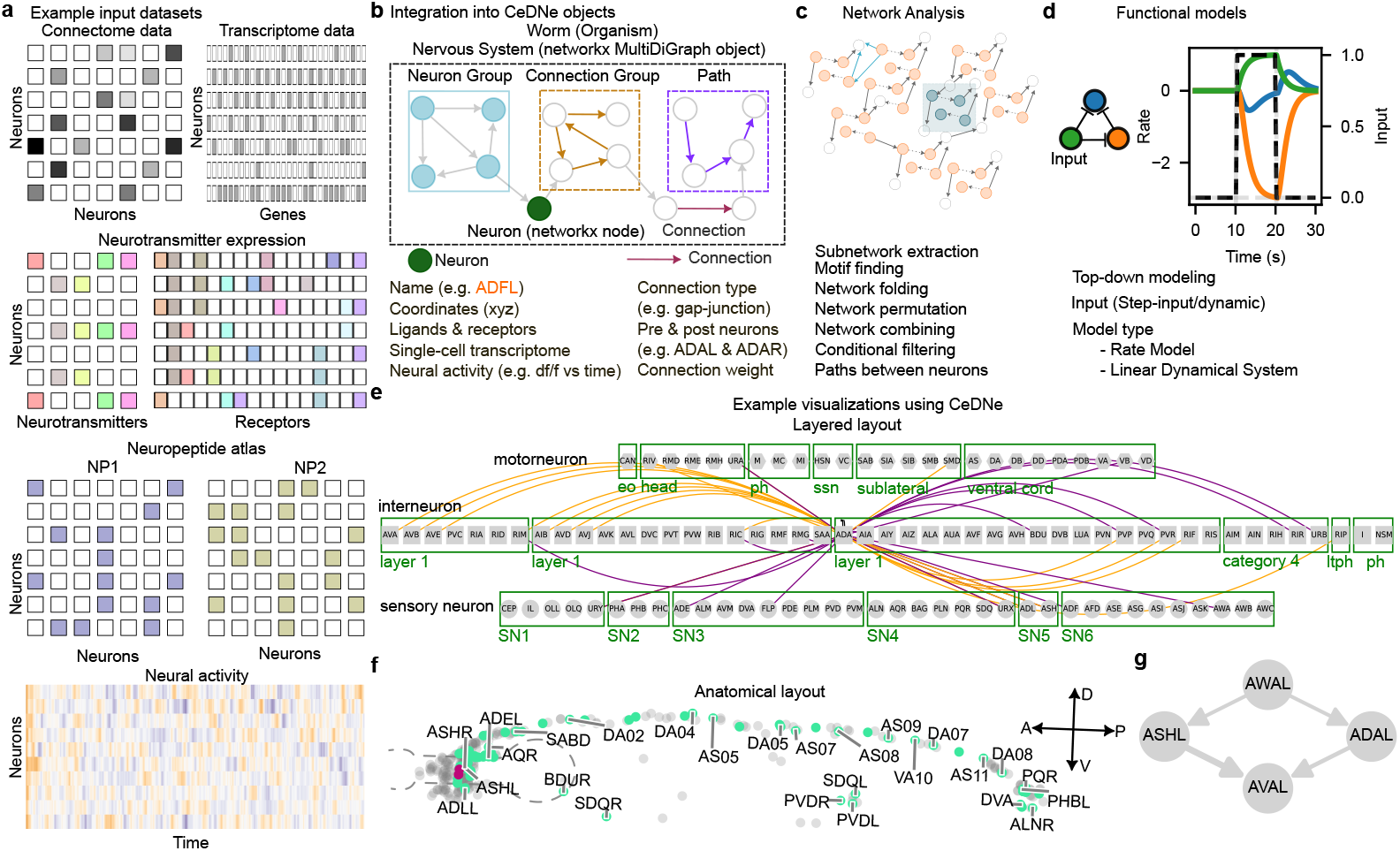
Schematic of CeDNe (see details in **Methods**) showing data integration, analysis, modeling and visualization enabled by CeDNe. **a**. Examples of diverse input datasets accessible to CeDNe-anatomical connectome, transcriptome, neurotransmitter ligand and receptor expression, neuropeptidergic connections, neural activity (top to bottom). **b**. Core components of CeDNe that constitute the unified graph object: Neurons (green), Connections (magenta), Neuron Groups (light blue), Connection Groups (gold), Paths (purple), and Nervous System (black), each of which contains biological attributes. Example datasets that can be integrated for *Neuron* and *Connection* components are shown below the graph. Custom datasets can be added by users. **c**. Examples of several network analyses enabled by CeDNe (**Methods**). **d**. The data-integrated graph objects feed into the modeling tools (simulator and optimizer classes) that facilitate top-down dynamic modeling of the worm nervous system. An inhibitory feedforward loop circuit shown here (top left) with a step input (black) between 10-20s given to the green node and dynamics produced in this circuit (top right). Users can currently access a few modeling tools (bottom). **e-g**. Users can visualize integrated networks in several layouts. Examples include-Layered layout which shows the whole nervous system grouped by cell-types [14], here shown with highlighted incoming (purple) and outgoing (orange) connections to ADA in (e); Anatomical layout showing the relative positions of the neurons, with a network or single-neuronal properties highlighted, shown here with AVAL and AVAR (magenta) and their connections (in light green) with a subset of neurons along the anterior-posterior axis labeled (f); Shell layouts that can be used to visualized several degrees of paths between neurons, here shown using AWAL and AVAL with path lengths of up to two from AWAL to AVAL in (g).

### Structural and anatomical insights from the *C. elegans* network

Understanding how network topology influences function requires integrating connectivity and molecular data. In this section, we demonstrate how CeDNe constructs unified network representations, enabling visualization, discovery and analysis of motifs, hierarchies, and connectivity patterns.

#### Visualizing network layouts

CeDNe allows multiple types of visualizations to explore anatomical and networklevel properties. First, using *anatomical layout* (**Fig**. 1**f**), we show spatial positions of AVA neurons in two dimensional space with their synaptic partners highlighted [14] in the anterior-posterior dorso-ventral plane. CeDNe also supports filtering based on synaptic strength and other planes of visualization as shown for ADA neurons in (**Extended Data Fig**.4**a**)

Next, to visualize information flow, we can use the *layer layout* 1**e**) to separate sensory neurons, interneurons, and motor neurons into layers and subcategories [14] (**Methods**) and highlight connections to and from ADA (input connections in purple, output connections in orange) in the network as shown in **Extended Data Fig**. 4**b**. This layer layout suggests a role of ADA in integrating information from sensory neurons and interneurons and in signaling primarily to other diverse interneuron groups. We can deepen the analysis by creating a *shell layout*, in which we isolate the connections and connected neurons for ADAL and ADAR neurons (**Extended Data Fig. 4b**) into a sub-network and display chemical synapses and gap junctions separately. This can be done for other connection specifications (*e.g*. glutamatergic connections) and neuron parameters (*e.g*. neurons that express *nmr-1* ). Thus, CeDNe enables flexible and systematic display of circuit connectivity of specific neurons or neuron groups at brain-wide scale with molecular resolution of individual neurons to facilitate hypothesis generation for neuronal functions and network properties across several scales.

#### Custom network partitioning

CeDNe enables flexible network partitioning into biological groupings based on gene expression, or function. Additionally, network can be partitioned into custom hierarchical groupings for task-specific investigations. For example, edges can be filtered based on connection type (synaptic, gap-junction, neuropeptidergic interactions). This flexibility allows users to zoom in on functionally relevant modules, facilitating both high-level circuit organization analysis and detailed mechanistic studies. Second, different research questions may require grouping neurons at different or even several hierarchical levels into interpretable units. These two applications are further illustrated with examples below.

First, mechanistic analysis of circuits can require following the flow of information across the nervous system. In such cases, knowledge of both direct (path-length = 1) and indirect paths (path-length >1) between a pair of neurons can be useful. Here we show monosynaptic (path-length = 1) and disynaptic paths (path-length = 2) from AWCL to RIAL using a shell layout (**Extended Data Fig**. 4**c**). The direct connection and the several indirect connections form multiple feed-forward loop motifs connecting this pair of neurons.

Second, neurons can be grouped together based on predefined conditions, such as gene expression, functional importance, etc or by user-defined conditions. Group connections can be defined to exist if any two members of a pair of groups connects, or if all members connect. CeDNe facilitates users to define custom groups to view group connectivity to the rest of the network, and thus allows access to the network from several hierarchical levels. This allows flexibility to look at the network from simple interpretable modules for high-level analysis, and then zooming into the modules for detailed investigations.

#### Motif analysis and hierarchical chains in *C. elegans*

In addition to visualization-aided exploration of network properties, CeDNe also enables investigation of high-level connectivity structure of the worm nervous system. Here, we use the analysis on triad motifs as an example. Triad motifs are sub-networks of 3 nodes that have been extensively characterized in several biological networks [17, 20] These motifs are not merely structural subunits; they are often regarded as building blocks of complex networks since they can transform inputs through specific computations such as gain control, oscillations, or multi-stability [20] First, we characterize the presence of triad motifs in the *C. elegans* chemical synapse connectome by searching for triad motifs through the adult hermaphrodite dataset [14], **Methods**. We found that triad motifs are significantly under- or over-represented, different from randomized networks (**Methods**, **Extended Data Fig**. 1**a**). This finding suggests a potential functional role of triad motifs in the worm neural network. Next, we asked whether the direction of information flow in triad motifs aligned with the hierarchical flow of information from sensory to motor direction. Thus, for each identified triad motif, we counted the number of connections that pointed *forward* (e.g. sensory to motor), *backward* (e.g. interneuron to sensory) and *lateral* (e.g. sensory to sensory) **Methods**. We then calculated *Hierarchical Alignment* as the normalized difference of the fraction of connections that point *forward* or *lateral* and the connections that point *backward* **Extended Data Fig**. 1 **b**, **Methods**. We found that feed-forward loop motifs were maximally aligned from the sensory to motor direction. Furthermore, the other motifs that had slightly lesser alignment than the feed-forward loop motif were either subsets or slight modifications of the feed-forward loop motif **Extended Data Fig**. 1 **b**. This trend is visualized in 2**a** showing that the fraction of sensory neurons matched with the input nodes is the highest and is significantly decreased in the output nodes. The trend is reversed for motor neurons, while interneurons are similarly matched for all three nodes.

Feed-forward loops are ubiquitously present in the invertebrate and vertebrate nervous systems, and have characterized functions in signal propagation, temporal filtering, activity normalization, signal amplification [17, 21]. They have also been previously characterized as an important motif in the *C. elegans* chemical connectome [14, 22]. However, motifs have not been analyzed previously in conjunction with other properties of the nervous system. Thus, we further analyzed *simple hierarchical chains* of feed-forward loop motifs (**Fig**.2**b-f**). There are two ways of combining feedforward loops into *simple hierarchical chains*. First, a *sequential hierarchy*, formed by joining the output node of the first feed-forward loop to the input node of the next. Second, *intermediate node-chains*, formed by joining the intermediate node of the first feed-forward loop to the input node of the next **Fig**. 2**b**. Note that there are several other kinds of non-hierarchical and mixed hierarchical chains possible, but here we chose to focus on simple hierarchical chains to reduce combinatorial complexity and maintaining our focus on the property of unidirectional sensory-motor information flow. We then searched for various lengths of chained motifs in the worm synaptic connectome in CeDNe. We found that a chain of 2-4 feed-forward loops joined together was overrepresented in the worm nervous system, as compared to random networks generated by swapping edges between nodes with a maximum at chains of 3 feed-forward loops (FFLs) (**Fig**. 2**e**,**Extended Data Fig**. 2**a,b**, **Methods**) . Furthermore, feed-forward loop motifs chained in lengths of 2 - 4 motifs into sequential hierarchies arrangements were more prevalent in the worm connectome over intermediate node chains **Fig**. 2**e**, opposite to the trend in randomized networks. The set of all sequential hierarchies composed together covered 218/302 neurons and 1100/3709 connections. Together, these results demonstrate a previously uncharacterized highlevel wiring principle of the worm nervous system. To address its significance for sensory information transformation and propagation, we analyzed connectome edges that corresponded to the different edges in the sequential hierarchy chains of feedforward loop motifs. We discovered a strong directionality to these motifs that aligned with the flow of information from sensory to motor neurons across the nervous system **Fig**. 2**c,d**. Such strong directionality was not found in the intermediate node-chained feed-forward loops **Extended Data Fig**. 1c or when searched in the randomized connectomes **Extended Data Fig**. 1d. In summary, by combining neuron-type annotations and the *C. elegans* connectome into a CeDNe network, we found a high-level organizational principle of information flow whereby a sequential hierarchy chain of feed-forward loop motifs aligns with the sensory-motor information transmission in the *C. elegans* nervous system. Although it is known that there is broad sensory to motor flow of information in the *C. elegans* nervous system ([13–15]), the enrichment of such chained hyper-motif structures is neither obvious, nor guaranteed. Nature’s preference of such structures suggests potential architectural or functional roles of these circuits in the *C. elegans* nervous system.

**Fig. 2.**
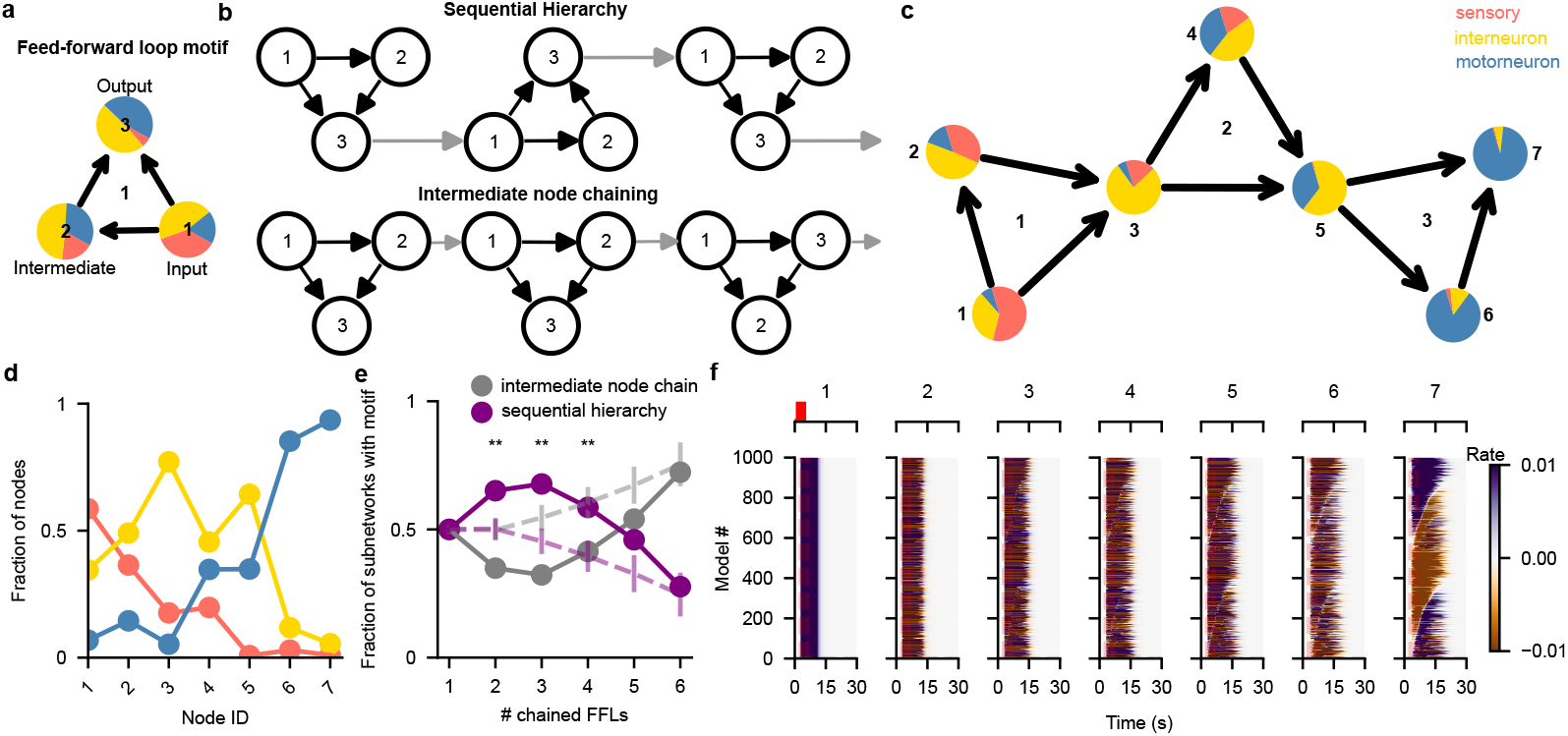
Chains of feed-forward loop motifs are prevalent in the worm nervous system and align to the sensory-motor flow of information. **a**. Feed-forward loop motif with input, intermediate and output nodes labeled and shown as pie-charts with fraction of sensory neurons (orange), interneurons (yellow) and motorneurons (blue) matched in the connectome as slices. Note the change in the fractions of sensory neurons, interneurons and motor neurons in the direction of information flow in the motif (1 −→ 3). **b**. Two ways of chaining feed-forward loops into simple hierarchical chains: sequential hierarchy (above) with the output node chained with the input node and intermediate node chains (below) with the intermediate node chained with the input node. **c**. A sequential hierarchy of length 3 with individual feed-forward loops labeled in the center, and nodes labeled outside. Fraction of neuron types matched for each node (pie charts same as **a**) show a strong alignment of information flow in these chains with the sensory to motor direction when matched to the chemical connectome. **d** The phasic change in fraction of sensory neurons, interneurons and motor neurons over the depth of the nodes in a length-3 sequential hierarchy. **e**. Fraction of sequential hierarchies (purple) in contrast with intermediate node chains (gray) of different lengths matched in real (solid lines) and randomized (dotted lines) networks. **f**. Percolation of a step input to the input node over the depth of nodes in a length-3 sequential hierarchy. Rows represent a different set of weights sampled for the sequential hierarchy and are sorted by the onset time for the output (7) node. Top inset for each shows inputs, depicting a 2 second step-input only provided to the first node (1) (in red).

### Sequential hierarchies can function to mediate temporal patterning of neural activity

We then wanted to test out how sequential hierarchies can transform inputs. Given the strongly directional nature of this architecture, its alignment to the sensory-motor axis, and its relative prevalence, we wanted to see how incoming inputs get transformed over the different nodes in this circuit. We used rate models to simulate these hierarchies and gave step input to only the input node (1) of the hierarchy (**Methods**). First, we fixed the gains and time constants of the nodes in the hierarchy to focus on the connections between the nodes. Then, we generated 1000 sets of sequential hierarchies by randomly assigning weight sets between -1 to 1 each to the nodes for each circuit (**Methods, Table** 1) to give us a distribution of input transformations through the circuit. We then gave each circuit an input pulse of 2 seconds, simulated this model using the simulator module in CeDNe for 30 seconds, and recorded the transformation through the different nodes. We found that the different weight sets between the nodes strongly modulated the onset and sustenance of the input through different nodes in the hierarchy (**Fig.2f**). Together, this tuning the onset and sustain of the output by weights between the nodes results in flexible temporal patterning of the input at the output nodes. This tuning results from the push-and-pull dynamics at the convergence of inputs at the intersecting nodes 3 and 5. If the weights of the incoming connections are in the same direction, they sum up and push the output of the node in one direction, but an opposing set of weights cancel the input transformations out, leading to a delay in the output from the node. Over the course of multiple loops, this can generate more complex temporal patterns at the output. This feature extends the gain control and temporal filtering attributes of individual feedforward loop motifs [17] in the sequential hierarchies to give rise to specific temporal patterns of activation in the output nodes (**Fig.2f**).

### Functional integration of neurotransmitters and gene expression

CeDNe enables the integration of multi-modal data with the anatomical connectome. Here, we extract data from the *C. elegans* neurotransmitter atlas gathered from reporter lines containing fluorescent markers inserted into endogenous loci [5], the expression of receptor genes at the highest threshold from CENGEN [2], the ligand-receptor map that matches the neurotransmitter ligands to their receptors [4], and map all of these expression profiles to the anatomical chemical connectome [14]. Together, these allow us to predict the type of neurotransmitters used by synaptic connections. We assign a connection between a pair of neurons a putative neurotransmitter if 1) There is a chemical synapse between the neurons based on the anatomical connectome, 2) The presynaptic neuron expresses the neurotransmitter ligand, 3) The postsynaptic neuron expresses the receptor for the corresponding presynaptic neurotransmitter. Based on this, we predict unique putative neurotransmitter types for 3709 chemical synapses. 567/3709 ( 15.2%) synapses could not be annotated due to a lack of matches between the set of neurotransmitters and receptors for the connection, 3070/3709 (82.7%) had a unique putative neurotransmitter and 72/3709 (1.9%) had two putative neurotransmitter types for the connection. In terms of neurotransmitter-receptor pairs, other than the unannotated 15.2% mentioned above, another 2831/3709 (76.3%) had 1-8 putative neurotransmitter-receptor pairs and the remaining 311/3709 (8.3 %) had more than 8 annotations **Extended Data Fig**. 3**a**. Multiple annotations result from the expression of more than one type of neurotransmitter and several receptor subtypes within neurons. In **Fig**. 3**a** we extend the ADA example and show predictions for neurotransmitter types for its synaptic connections. We can also isolate specific neurotransmitter-receptor pair annotations and create a complete graph for that neurotransmitter type. In **Fig**. 3**b** we show the complete putative glutamatergic connectome. We note that while previous studies have used older datasets to make similar predictions [23], we use our predictions for the annotated connectome built using the more recent datasets and integrate it with network properties to discover multi-scale and functional patterns shown below.

Next, we analyze the large scale connectivity structure of chemical connectomes and provide uniquely annotated putative neurotransmitter types for all connections grouped by neuron classes as defined in [14] (**Fig**. 3**b**). Here, a pair of groups is connected to each other if *any* member is connected across the groups. Here we observe in the “circular layout”, several features of the partitioning of the nervous system by neurotransmitter-receptor pairs: glutamatergic connections are never involved in feedback from the motorneurons, although they can be involved in both feedback from interneurons and also feed-forward inputs to both interneurons and motorneurons (**Fig**. 3**b**). Similarly, we can see the considerable local and feed-forward structure in the dopaminergic connectome.

**Fig. 3.**
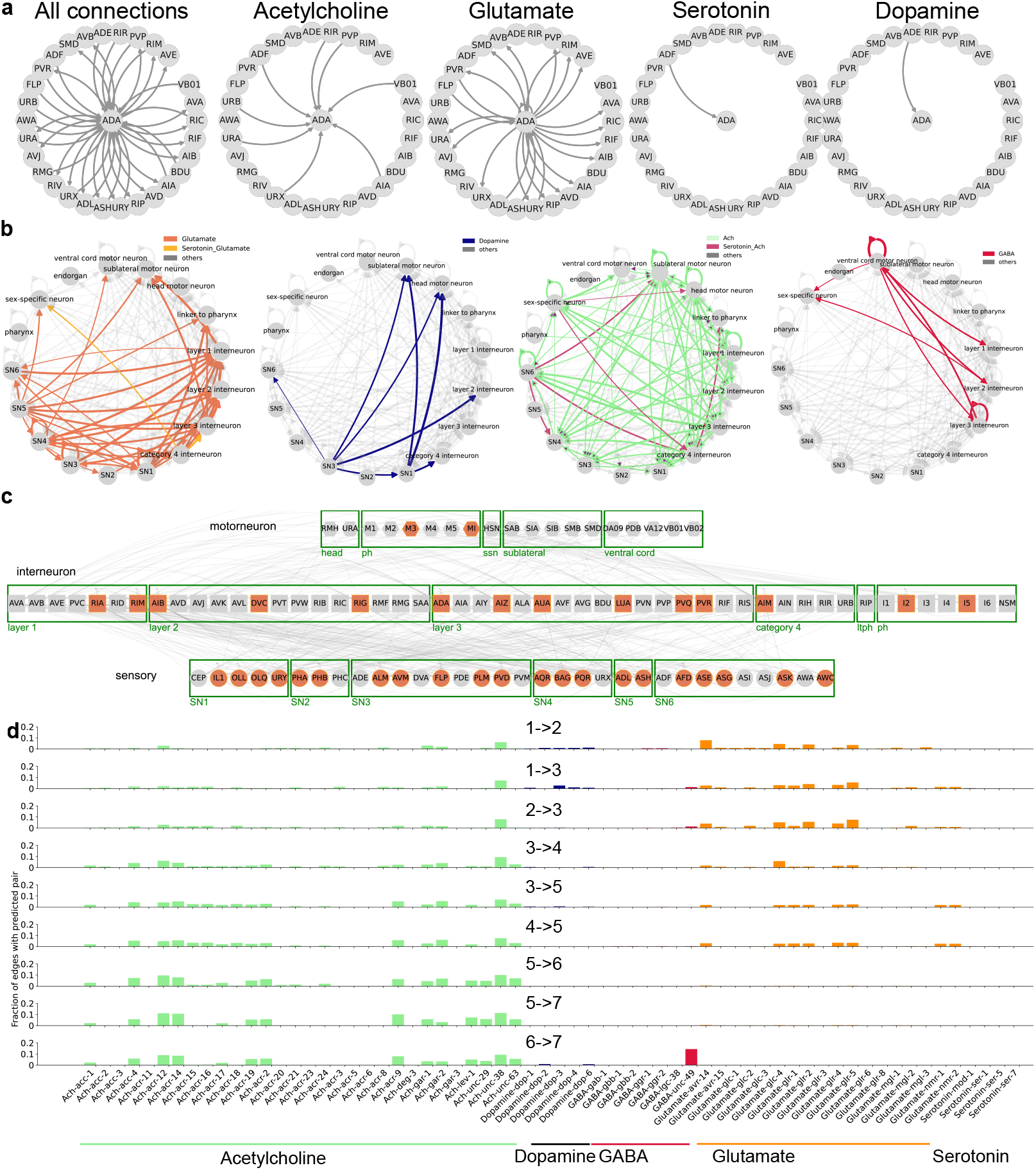
Neurotransmitter annotations for the connectome reveal a partitioning of the nervous system over chains of feed-forward loop motifs. **a**. Neurotransmitter annotations for synaptic connections to and from ADA. There is no GABA or tyramine connection for this neuron. **b**. Large scale connectivity grouped by neuron class according to [14] and colored according to putative neurotransmitter-receptor pairs, in order: Glutamate (left), Dopamine, Acetylcholine and GABA (right). In case the group had connections with more than one predicted neurotransmitter, they were colored differently. **c** The complete putative glutamate connectome with neurons that receive at least one glutamatergic connection (participating connections and neurons in gray, neurons that also release glutamate in orange). Other neuron classes were omitted from the graph. **d**. Distribution of neurotransmitter ligand and receptor pair annotations for edges in the chemical connectome matched by the sequential hierarchy in **Fig**. 2**c**. Cholinergic connections are marked in green, dopaminergic in blue, GABAergic in red and glutamatergic in orange. Other pairs were not significantly represented.

Finally, we characterize the feed-forward loop motif (**Extended Data Fig**. 3**d**) and the sequential hierarchy motif (**Fig**. 2**c**) using the neurotransmitter-receptor pairs added above (**Fig**. 3**d**). Strikingly, we found that the connections gradually transitioned from glutamatergic to cholinergic across the connections starting from the input node 1 to the output node 7 (**Fig**. 3**d**). Furthermore, there is an especially large contribution from the neurotransmitter GABA - *unc-49* receptor pairs to the final connection between the intermediate node and the output node. *unc-49*, known to encode a GABA-A receptor subunit is essential for the inhibition of motorneurons and plays a critical role generating locomotion [24]. The strong enrichment of the last node in motorneurons (**Fig**.2**c**) and consistently, the enrichment of unc-49 ligand-receptor connection at the final connection to the output node suggests that sequential hierarchies may have functional role in the unidirectional propagation and temporal transformation of input from the sensory to motor circuits. In summary, by combining several sources of data, we generated putative neurotransmitter connectomes for different neurotransmitters and found a detailed-level partitioning of the nervous system. We showed that a *sequential hierarchy* constructed by chained feedforward loop motifs use neurotransmitters in a structured direction aligned with the sensory-motor information transmission in the *C. elegans* nervous system.

### Integrating and visualizing neuropeptide-receptor types and gene expression data

In parallel with neurotransmission via chemical synapses and gap junctions, neurons also communicate using a rich repertoire of neuropeptides that are released into synaptic or extrasynaptic regions from dense-core vesicles to act on their receptors locally or over a distance. Thus, neuropeptide-mediated intercellular signals add layers of complexity and expand the spatial and temporal scales of neuronal communication. We integrated the neuropeptide-receptor maps from [6] and investigated enrichment of neuropeptides in the connections between pharyngeal neurons to illustrate a usecase of CeDNe in hypothesis generation. We found that several neuropeptidergic connections in the pharyngeal circuit were enriched (**Fig**.4b), making them suitable candidates for functionally important roles within the pharynx, such as modulation of pumping or feeding-related behaviors.

**Fig. 4.**
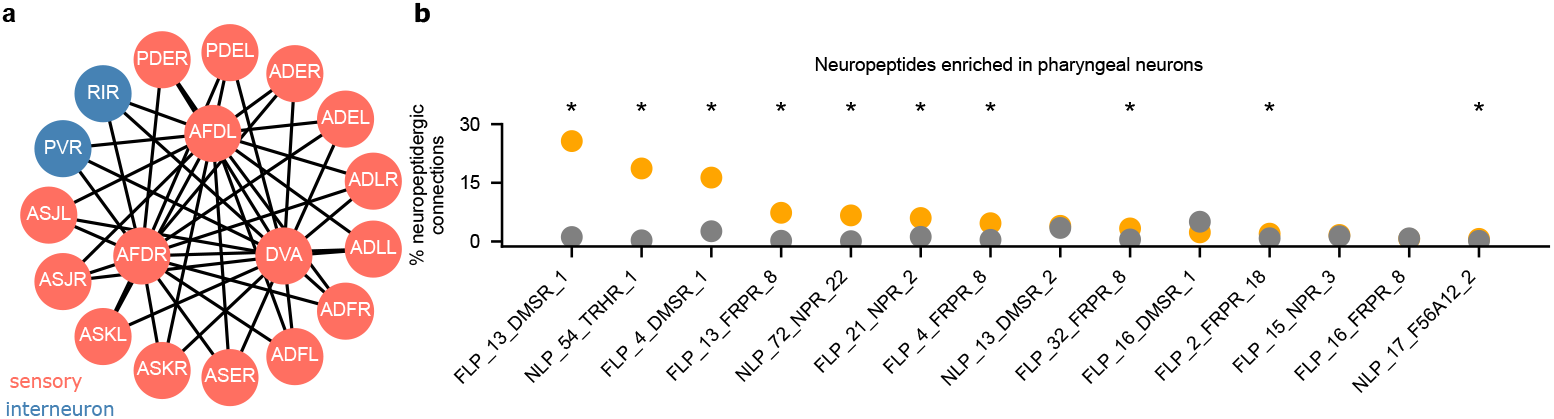
Integrating the neuropeptidergic connectome into CeDNe to find circuit enriched neuropeptides. **a**. The NTC-1 to NTR-1 neuropeptide connectivity map using long-range model [6]. The inner shell represents the neurons that express NTC-1 (senders) and the outer shell the neurons that express NTR-1 (receivers). Node colors indicate neuron types (orange - sensory, and blue - interneurons) **b**. Enrichment analysis of neuropeptidergic connections between pharyngeal neurons. Asterisks indicate significant difference between neuronal subset (in orange) and the rest of the network (in grey) (z-test, p<0.05, Benjamini-Yekutieli false-discovery rate correction).

Similarly, the expression pattern of a specific gene of interest in the nervous system can be useful for several experiments. In **Extended Data Fig**.6**a**, we show the expression pattern of two GPCRs sre-1 and sre-2 in purple and green respectively. The overlapping neurons are shown in dual colored circles. While this example takes a single pair of genes, CeDNe supports this operation for any number of genes together.

### Integrating functional calcium imaging data

CeDNe allows integration of whole-brain imaging datasets in order to contextualize neural activity data within the scaffold of other relevant information. CeDNe’s graph structure, integration and analysis functions allow users to store imaging data in the CeDNe structure, which can be useful for quickly aligning data across datasets, folding and averaging classes of neurons, such as left-right neurons of a given class and testing out patterns between neural activity correlations and anatomical connectivity. [12] shows a first use of CeDNe with calcium imaging data. Here all calcium data was stored in CeDNe’s relational data structure, allowing access to neural data from a hierarchical data structure and enabling analysis, visualization and simulation from the same unified data objects. This ensures reproducibility across the different aspects of the analysis as all data transformations can be tracked back to the same data objects. For example, in Figures 1,2 and 5 of [12], the same unified data structure is used to store raw imaging data (shown in fig 1d and e), for visualization, analysis and simulation. Further, in Figure 1b of [12], the authors used promoter lines that labeled multiple neurons with partial overlaps. Using the anatomical layout the authors show anatomical positions of the labeled neurons belonging to different lines colored by their promoter types, showing the expression overlaps between the promoters.

Additional tutorials on integration of functional data will be made available on our Github page upon acceptance.

### Generalization to other organisms and extensions

We have assembled several other datasets for *C. elegans*: developmental connectomes for hermaphrodites, [15], male connectome [14] and neurotransmitter information [5]. A more complete list can be found on the online repository. We plan to extend the broader platform to other model organisms for which multiomic data at the cellular scale are available, such as *Drosophila melanogaster, Platynereis dumerilii, Pristionchus pacificus* etc. CeDNe supports integration of data in various formats including adjacency matrices, microsoft excel files and JSON files among others. We plan to extend support for other formats such as NWB over time.

Here we show 2 examples of integrating other datasets in CeDNe. First, we analyzed whether the sequential hierarchies and their properties are conserved in the male connectome. We integrated the male *C. elegans* connectome and their annotations, and repeated the process from **Fig**. 2**a-e** by chaining different lengths of feed-forward loop motifs and matching the pattern in the worm chemical connectome. We found that length-3 sequential hierarchies were uniquely over-represented in the male connectome, a trend similar to hermaphrodites **Extended Data Fig. 5a**. Furthermore, we found that they were also strongly aligned in the sensory to motor direction **Fig**. 5**a**. Then we integrated the neurotransmitter information for males from [5] and made predictions for neurotransmitter-receptor pairs for chemical connectome edges. Again, we discovered strikingly similar distribution of neurotransmitter receptors that matched for the different edges of the sequential hierarchy suggesting that these high-level structural features are conserved for the *C. elegans* nervous system.

**Fig. 5.**
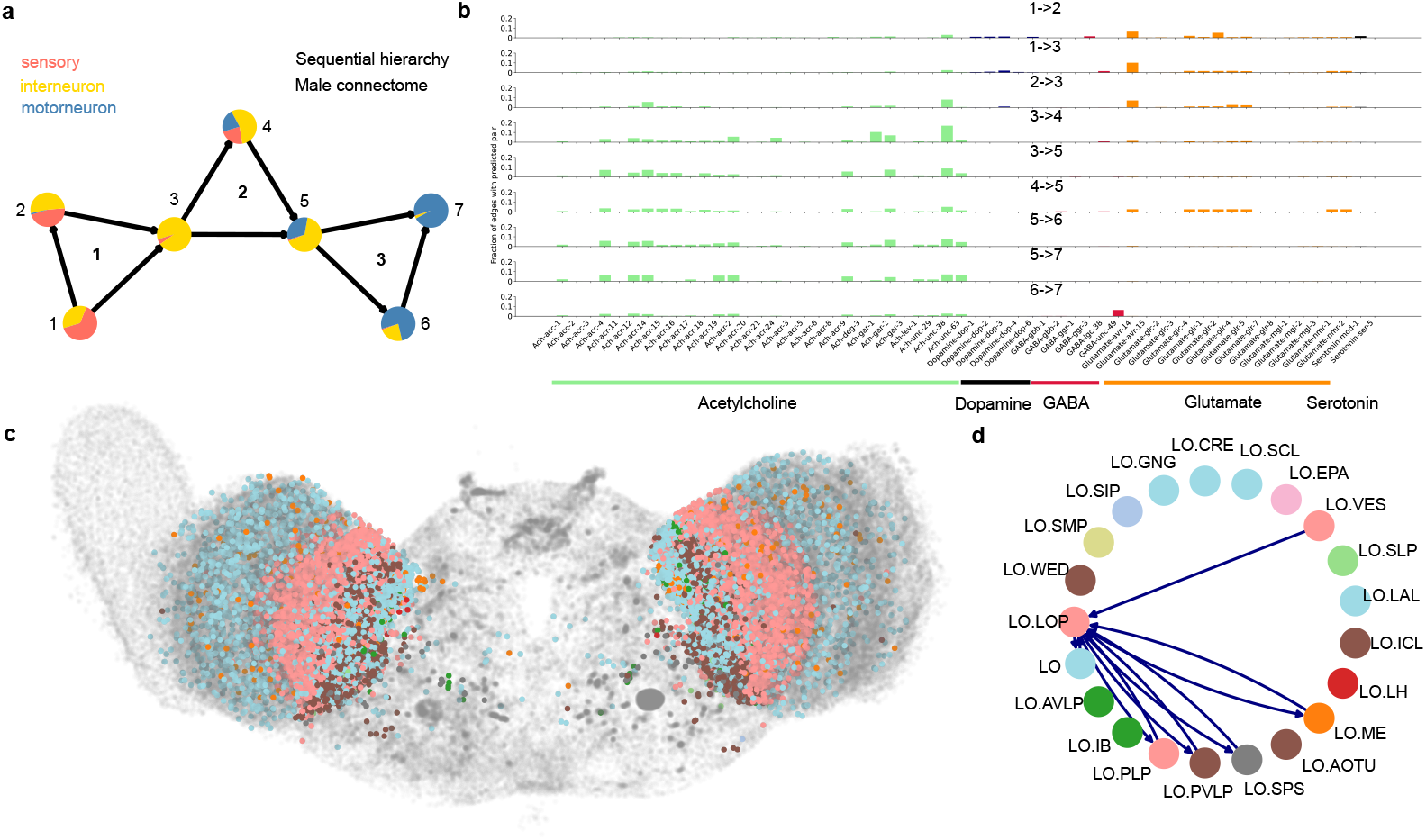
CeDNe extensions to other species or sexes using the same classes and pipelines. **a**. Sequential hierarchy in the male connectome showing the same pattern of hierarchical alignment as the hermaphrodite. Node colors represent neuron types (orange-sensory neurons, yellow-interneuron, blue-motorneuron). **b**. Neurotransmitter partitioning in the male connectome shows a similar progression from glutamatergic to cholinergic putative assignments in the direction from sensory neurons to motor neurons. **c**. Neurons from the EM reconstruction of the fly brain shown as a scatter plot, colored by different subtypes within the Lobula. [25, 26] **d**. Neurons from **c** grouped by their subtypes and their connections aggregated (colors same as **c**). Dark blue edges represent dopaminergic connections between Lobula Plate (LOP) subclass and the rest of the Lobula neuron subclasses.

Next, we integrated the recently published adult *D. melanogaster* connectome [25, 26] in CeDNe **Fig**. 5**c**. Here we show a map of the fly brain with several neuronal subtypes in the Lobula. The fly connectome already has annotations for anatomical positions of neurons, neuron classes and neurotransmitter types for synaptic connections. We chose the optic lobe to focus on due to its anatomically compartmentalized classes for ease of visualization. We first *folded* neurons of a given neuron class within the Lobula into class nodes (*e.g*. LO.LOP, LO.VES, *etc*. (**Fig**. 5**c,d**)), and assigned a different color for each class. Then, we plotted in anatomical layout the fly brain colored by neuron classes 5**c**)). We then used CeDNe to select connections that were labeled dopaminergic between one of the classes (LO.LOP) and the rest of the subclasses in the class 5**d**)) demonstrating flexibility in CeDNe’s application to other datasets.

### Tutorials, website and web interface for easy navigation

In order to make these tools available to a wide audience, we provide several tutorials and example scripts in the form of Jupyter notebooks that demonstrate how to do several operations in CeDNe. These are available at our Github repository at https://github.com/sahilm89/CeDNe as online executable Jupyter notebooks using Binder [27]. Finally, advanced users and developers are invited to contribute to CeDNe Github repository for issue tracking and collaborative feature addition and model building.

## Discussion

Over the past decade, several key technologies such as high-throughput sequencing, molecular-biology tools, optogenetics, brain-wide calcium imaging, and machine vision have matured. This has precipitated a rapid increase in organism-level neuroscience data, especially for *C. elegans*. The field has reached a point where it has become practical to meet the demand for organism-level models that integrate several independent sources of information to address systems-level questions that depend on bridging knowledge and insights distilled from different omics-scale datasets [16, 28]. To address this demand, here we present a unifying framework, CeDNe, whose goal is to integrate the multi-modal datasets available for the worm and build an evolving model of the worm nervous system. As examples to demonstrate the unique strengths of CeDNe, we identified overrepresented feed-forward loop chains aligned with the direction of sensory-motor processing, demonstrated a hierarchical neurotransmitter organization and showed its generality using the male connectome. The observed hierarchy of feed-forward loops suggests a structured propagation of information through the *C. elegans* nervous system. Moreover, the predicted neurotransmitter assignments offer testable hypotheses for circuit function, guiding future experiments in optogenetics and functional imaging. While existing approaches such as WormAtlas provide static connectivity maps, CeDNe enables network-wide motif discovery, functional prediction, and cross-modal integration. Unlike bottom-up biophysical models, which require detailed electrophysiological parameters, CeDNe offers a scalable top-down modeling approach [12], making it useful for broad exploratory analyses and hypothesis generation.

There are several ways in which experimentalists can immediately make use of CeDNe’s approach to guide experiments.

### Ranking potential perturbation targets

Experimentalists often need to search for elements of a pathway- such as neuropeptidergic signaling, or gene expressions, by disrupting several alternative candidates in a screen. CeDNe can help reduce the search space by making predictions about enrichment from existing datasets. For example, neuropeptides enriched in specific subnetworks that can be preferentially targeted for disruption (**Fig**. 4**b**).

### Visualizing patterns on the connectome

Using several layouts, CeDNe allows visualization of intersections of various datasets, *e.g*. merging gene expressions and anatomical positions to give anatomical expression maps for multiple genes (**Extended Data Fig**. 6a).

### Data analysis

CeDNe allows storage of data in a graph structure that immediately puts imaging data in the context of the connectome. Further, CeDNe’s hierarchical structure and analysis tools allow for quick hypothesis testing. For example, by integrating the results from the brain-wide functional imaging of naive and trained worms in an aversive learning paradigm, we examined the anatomical organization of the neurons carrying information of contexts and learning in the wiring diagram, as shown in Figure 1g in [12].

## Key features

CeDNe has been built with the aim to integrate, intersect and visualize information sources, in order to build evolving models of the *C. elegans* neuronal network. Below we lay out CeDNe’s key features.

### Evolving models framework

Top-down phenomenological model presented here helps discover interactions between neuronal activity without necessarily having an accurate biophysical interpretation of its generative microscopic variables such as ion-channel distributions, membrane resistances, etc. We envision this tool’s core function as building top-down organism-level, evolving models, which should iteratively improve as more organism-level data gets integrated into this framework, thus building an increasingly annotated connectome [29].

### Integration of omics and data synergy

CeDNe provides a ground for integrating connectomic, transcriptomic, and other omic datasets by organizing information together, enabling cross referencing, and providing tools to find intersections. There are excellent tools already available that focus on individual aspects of these datasets. In comparison, CeDNe aims to scaffold these datasets into one cohesive framework. This system allows discovery of insights contained within the relationships between independent sources of data. For example, CeDNe allows users to identify neurons expressing a specific gene and at the same time examine their connectivity patterns across the network.

### Contextualization

Due to the open and collaborative nature of *C. elegans* research community, many large neuroscience datasets are published in open-access formats. Additionally, there is a strong push for open data in neuroscience both from NIH guidelines and from the scientific community. Data standardization formats [30] are increasingly being adopted by neuroscientist in service of this goal [31]. Taking another step in the direction of unification, integrative and accessible *evolving models* have the potential to enhance collaboration between labs working on different subsystems, and more directly use existing knowledge as context for their work. Importantly, integrated models must comply with the data sharing and licensing guidelines of the strictest license among the datasets that they are built on. Lastly, these models can guide experiment design by reducing both the collection of redundant data and the space of potential experiments.

### Analysis

The library underlying CeDNe is networkx, which is built for advanced graph-theory algorithms. This allows flexibility and scalability to CeDNe for integrating both future worm datasets and also other organisms with larger networks. We have leveraged this library to build neuroscience-relevant features to enable flexible data-analysis and model building. For example, drawing summary graphs based on custom grouping of neurons according to neurotransmitter identity, promoter expression, help dissect the network into interpretable subunits relevant to individual research questions. Similarly, analyzing chemical, gap-junction and neuropeptide networks independently as well as in union can help identify their different roles in generating network activity or behavior.

### Visualization

CeDNe allows users to flexibly draw data-enriched graphs. Several different layouts customized from networkx are provided. Users also have the flexibility of exporting graphs to other graph drawing libraries using popular graph formats. While there are existing tools that allow visualization of edges labeled by neurotransmitter identity and gene expression (www.nemanode.org, wormwiring.org), to our knowledge, this is the first such effort towards a systematic unification of the various datasets available for the worm with the purpose of integrative analysis and functional model building.

### Accessibility

CeDNe has been designed for users with varying levels of programming expertise. The Python library has been documented and several tutorials are provided to enable ease of access. There is additionally a web interface that can be used to visualize omic-level properties.

#### Challenges

Integrating large datasets comes with several challenges that emerge from 4 main categories of necessary assumptions.

### Stereotypy and stability

Given our current state of knowledge, anatomical connectomes and the transcriptomes are assumed to be reasonably stereotypical and relatively stable across individual animals under similar conditions.

### Symmetry within neuron classes

Different datasets acquired at the whole organism level are not all at the same level of granularity. For example, the singlecell transcriptomic data [2] is categorized by cell-types that group left-right pairs or dorsal-ventral pairs, while the anatomical connectome separates out individual neurons. We impute the transcriptomic (and other) datasets available for different neuronal groups to match the more granular dataset of the anatomical connectome with the assumption that the transcriptome in the same type of neurons is similar. Similarly, the datasets were not always obtained from the same developmental stage. For example, the single-cell transcriptomic data is from the last larval stage, L4, of the worm, while the connectomic data is from the first-day adults ([14] or L1 - L3 stage larvae ([15]).

### Consistency across datasets

Individual organism-level datasets may not always have the same degree of coverage of the nervous system. This could happen, for example, due to technical limitation, partly unusable data or by experiment design. Thus, different datasets may have to be patched together to generate a complete dataset. Similarly, there can be large variability in functional datasets across labs. This can be due to differences in conditions or specific experimental methods.

Thus the integration of the currently available datasets is imperfect. We therefore employ several methods in order to work around these challenges. We implemented network folding and data aggregation methods in CeDNe (Methods) and use consensus models, whenever possible for higher robustness. Further, CeDNe allows integration at the level of the coarsest dataset or imputing low granularity dataset across classes to integrate at finer levels, so that users have control over their assumptions.

#### Extensibility

It is likely that with continued development of high-throughput organism-wide data acquisition technologies, including cryo-electron microscopy, single-cell RNA sequencing and functional imaging, more datasets of connectivities, expression patterns of ion-channels, neurotransmitter, neuropeptides and their receptors, and gap-junctions will continue to be acquired. In addition, we also expect to see more contingent gene expression data under different states of the organism, e.g. naive and trained animals upon learning [32]. Our tool can continuously integrate these and other future datasets to allow for comparison with existing data using the framework of multi-relational graphs.

#### Position in the ecosystem of neuroscience tools

Excellent platforms for the *bottom-up* biophysical modeling of the nervous system of *C. elegans* [7, 18, 19, 33–41] have been built that aim to simulate the worm neural activity and behavior using biophysical models and experimentally constrained parameters or top-down abstract models [42, 43], but without integrating multi-modal datasets available for *C. elegans*. CeDNe builds from the *top-down* direction of finding appropriate coarse-grained models from structural and functional data . We envision that these approaches will complement each other in modeling neural activity and behavior (**Extended Data Fig**. 7a).

Furthermore, nemanode [15] and wormweb.org are intuitive and descriptive tools for visualizing network properties. CeDNe does not aim to be a substitute for these existing tools. While CeDNe enables such visualizations, it is focused on cross-omic integration, analysis and visualization of the nervous system.

Overall, we envision that CeDNe will benefit the worm community at large in its ongoing efforts to integrate the systematic characterization of the development, structure, and function of the nervous system. Further, CeDNe’s core infrastructure is flexible and can be adapted to other organisms with a similar scale of datasets (as shown for the fly in (**Fig**. 5**c,d**), or for brain regions for which cellular-level multi-modal data is available, such as zebrafish and mouse.

## Methods

### Platform Architecture

The general design principle of this platform is five-fold. First, **data-driven**. The library organizes available neural data into a multi-relational graph structure (directed multi-edge network) that resembles the hierarchy of organization of the nervous system. Second, **modular**. It enables modular analysis of structural properties of different kinds of graphs of the nervous system. Information can travel down the nervous system through various routes, including gap-junctions, synaptic transmission by different neurotransmitters, non-synaptic transmission mediated by secreted peptides or proteins with their receptors. Each of these can impose a connectivity graph on the neurons, which can be informative to analyze both independently and in relation to each other. Third, **unifying and synthesizing**. Whole-nervous system level datasets are rich resources that could provide even richer information about the nervous system when analyzed together. This can be useful for contextualizing newly acquired data with the existing knowledge and assist in scaffolding the data as more whole-organism level data are acquired in the future. The interaction between different connectivity graphs and sensory stimuli direct the flow of information and activity across the nervous system. Thus, new temporal data such as neural activity and behavior could be integrated within the context of the structural properties of the underlying graphs, which can enable the generation of mechanistic hypotheses. Fourth, **extensibility**, to allow for new kinds of data to be integrated into this setup, and to enable new tools to be added and existing tools connected into this framework. And fifth, **adaptability**- CeDNe can be adapted for use for different model organisms which have multi-modal cellular-level datasets at the whole-brain scale, such as fruitfly and zebrafish; to partially available multimodal cellular-level datasets, such as in the mouse; and to other organisms who have datasets that are not cellular level, but can be cast into multi-relational graphs, such as humans.

Several of the datasets referred in this manuscript have their own fully functional and independent web interfaces, where users can access these resources. Thus, the purpose of this library is not as a place to access these existing resources. Rather, it is to build evolving models that can integrate these and future disparate sources of information about the nervous system, with cells and their connections as the base of the information structure.

In order to achieve this, we built an object-oriented platform that organizes neuronal data into graphs. At the top of this hierarchy is the class *Worm* (W) with attributes ‘Name’, and ‘Conditions’ to label the worm and experimental conditions that the worm could be in, e.g. Worm *glr-1_mutant_001* with condition *Control*. An instance of the *Worm* class contains a *Nervous System* object and a *Behavior* object. This class can also contain other user-specific properties or other temporal data.

The *Nervous System* is the workhorse class that is built on top of the networkx MultiDiGraph class that is a directed network which can support multiple edges between nodes **Fig**. 1**b**. Here, class *Neurons* are the nodes, which store several preexisting properties, such as single-cell transcriptome, ligands, receptors, gap-junction molecule identities, etc. Edges contain data about the connections between neurons-synaptic, gap-junction, neuropeptidergic, etc. The library also allows users to add arbitrary new properties to objects at each level.

Thus, the Python library presented here holds different datasets from the *C. elegans* nervous system in the form of a collection of graph structures. The library uses a few main custom classes: Neuron, NeuronGroup, Connection (inter-node interactions), ConnectionGroup, and Network (collection of nodes & edges) to represent neurons and collection of neurons, cross-neuronal interactions and their collection, and the nervous system respectively. Each class here is also a data-container holding both data structures and subroutines. These containers hold relevant parameters such as neuron labels, neurotransmitters, receptors, time-series data, etc **Fig**. 1**b**. The library is built keeping a modular structure in mind, and new data and subroutines can be easily added on top of this underlying multiedge-directed graph structure. Models can currently be saved as Python pickle files, so that all custom model parameters are saved within the model.

The library has ample documentation, and also includes several utility functions that help users in making efficient use of the library by using single-line functions to load different freely available datasets into a model, plot different kinds of graph visualizations, and analyze data.

### Graph Processing and analysis

#### Motif search

Motif search in CeDNe was performed using VF2 based Directed Graph isomorphism method in NetworkX [44].

#### Generating random networks

Random networks were generated by swapping edges of the network sequentially for about 30000 times (number of edges × log(number of edges)). This maintains both the in and out-degree of the nodes in the network, while dismantling the higher-order connectivity structure of the network.

#### Hierarchical Alignment

Hierarchical alignment (H) was calculated by counting directed connections *C*_*i*→*j*_ follows:

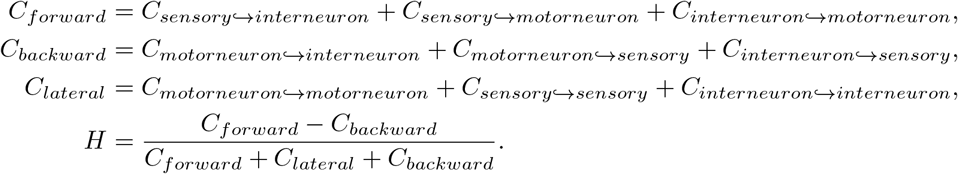

#### Network folding

Network folding involves merging together nodes or edges of a network based on a grouping criterion. We used CeDNe to *fold* the networks to get node-class representations as new nodes using a folding dictionary. For example, for folding left and right neurons into a neuron-class node, we made a folding dictionary with the class-name as key and the neuron names as values. CeDNe then collects the edges (and associated data) from each neuron and merges them into the class node.

##### Subnetwork extraction

We extracted node-subnetworks and edge-subnetworks using CeDNe’s subnetworking functions. These build on top of networkx subgraph functions, which subsets a graph based on a list of nodes or edges, by also copying the data contained in the *Neurons* or *Connections* and returns a new CeDNe *NervousSystem*.

### Statistical tests and validation

One-sided z-test was performed for comparing real networks against randomized networks by testing the null hypothesis that 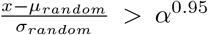, where *α*^0.95^ = 1.96, defined as the critical value for z-test at 0.95 significance. We corrected for False Discovery Rate using the Benjamini-Yekutieli procedure.

#### Modeling sequential hierarchies

We used a rate model to simulate the sequential hierarchy in CeDNe. CeDNe contains a *Model* class, which can be sub-classed to create custom models, both from scratch and from other networks in CeDNe. Currently, the default inbuilt model is a rate model, defined by class *RateModel* that implements the following model:

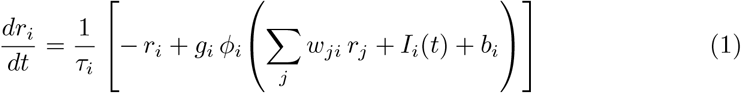

where
*r*_*i*_(*t*): firing rate of node i, *τ*_*i*_: membrane time constant, *g*_*i*_: gain, *w*_*ji*_: synaptic weight from node j to node i, *I*_*i*_(*t*): external (input) drive, *b*_*i*_: baseline term, *ϕ*_*i*_(*·*): activation function.

**Table 1.** Parameters used for the rate-model simulation of the sequential hierarchy.

| Parameter | Symbol / Value | Description |
| --- | --- | --- |
| Chain length | 3 | Number of motif units connected in sequence |
| Number of nodes | 7 | 3 nodes per motif with 2 intersection nodes |
| Input nodes | 1 | Nodes receiving external drive |
| Gains | $g_i = 10^{0.1} \approx 1.26$ | Uniform gain across all nodes |
| Baselines | $b_i = 0$ | Baseline firing rate offset for all nodes |
| Time constants | $\tau_i = 1$ | Integration time constant for all nodes |
| Weights | $w_{ij} \sim \mathcal{U}(-1, 1)$ | Random edge weights drawn per iteration |
| Iterations | $N_{\text{iter}} = 1000$ | Number of independent weight randomizations |
| Maximum time | $T_{\text{max}} = 30$ | Total simulation duration (seconds) |
| Time steps | $n_t = 1500$ | Number of discrete integration steps |
| Initial rates | $r_i(0) = 0$ | Initial activity of each node |
| Input onset | $t_{\text{start}} = 2$ seconds | Step input begins |
| Input offset | $t_{\text{end}} = 4$ seconds | Step input ends |
| Input amplitude | $I_1 = 1$ | Step magnitude of external drive to node 1 |

## Code availability

The code and documentation are available on our GitHub repository at https://github.com/sahilm89/CeDNe and will also be accessible via the website https://CeDNe.org

## Acknowledgments

Funding sources. Y.Z.is supported by NIH (NS115484, MH130064)

## Extended Data Figures

**Extended Data Fig. 1.**
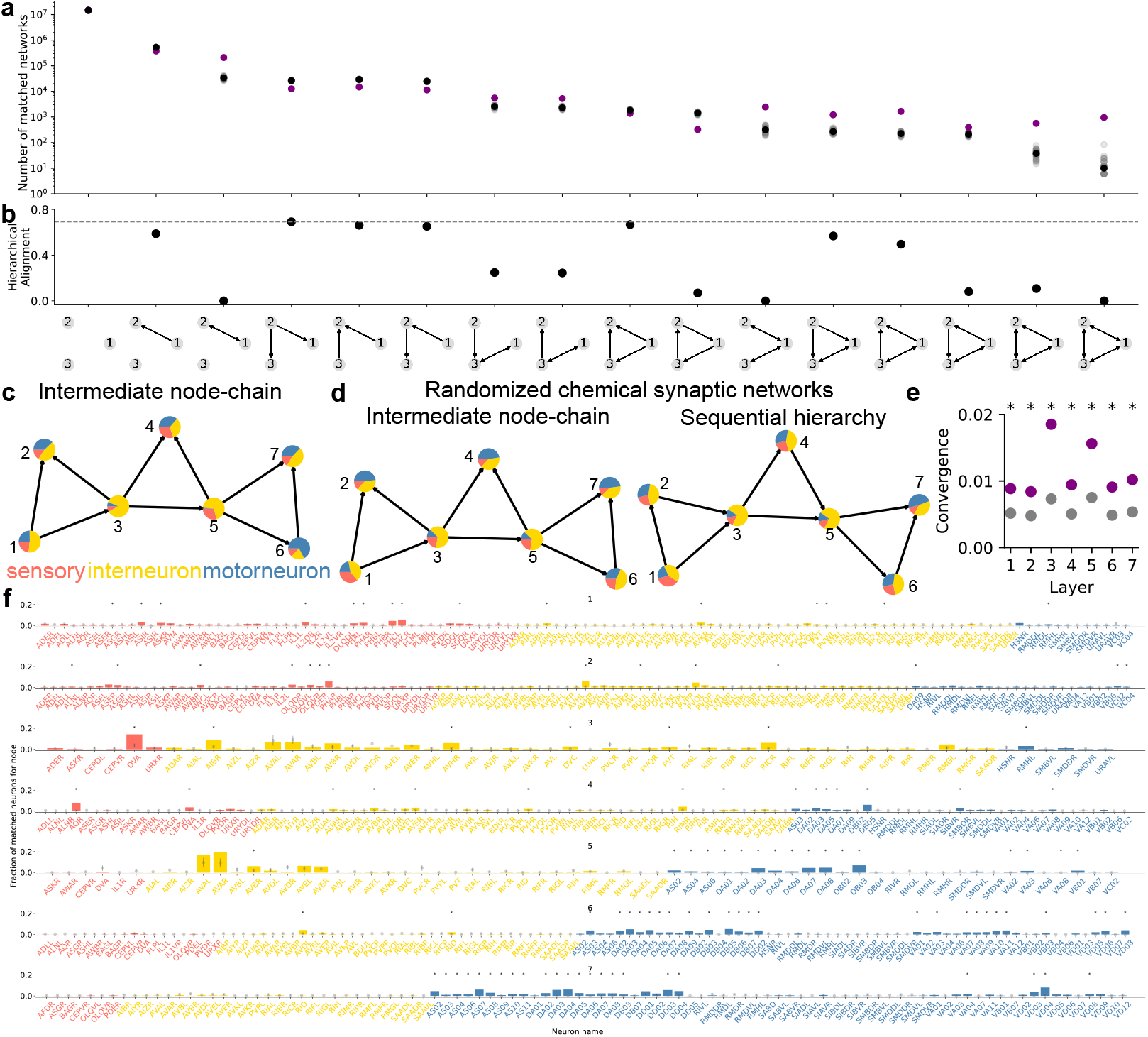
*C. elegans* hermaphrodite connectome shows high prevalence of circuit motifs that facilitate unidirectional sensory-motor directional flow. **a**. Motif search for various triad motifs in the *C. elegans* chemical connectome. All motif structures were significantly different in the real connectome (purple) when compared to the connectome randomized 50 times by edge-swapping (gray, average in black). **b** Hierarchical alignment for different motifs to the sensory-motor direction. Feed-forward loop (motif 9) had the strongest alignment to the sensorymotor direction. Other motifs with high alignments were either small modifications (motifs 12 and 13 (bidirectional edges), or subsets (motifs 2, 4, 5 and 6) of the feed-forward loop motif. **c**. Intermediate node-chains with fraction of sensory, interneurons and motor neurons matched for each node does not show a strong directionality as seen in **Fig. 2c. d**. Motif search for sequential hierarchies in randomized networks does not show the hierarchical alignment as in 2c, but intermediate node-chains (in c) do not show such dramatic differences. **e**. Layer-by-layer convergence onto a subset of neurons for a sequential hierarchy in the *C. elegans* connectome (purple) compared to 50 edge-swapped randomized connectomes (gray dots are mean and bars standard deviation, right-sided z-test *pvalue <* 0.05). **f**. Histogram of neurons in the *C. elegans* connectome that match for each node of the sequential hierarchy in 2c colored by neuron type, compared to 50 edge-swapped randomized connectomes (gray dots are mean and bars standard deviation, right-sided z-test *pvalue <* 0.05) with Benjamini-Yekutieli multiple comparisons correction.

**Extended Data Fig. 2.**
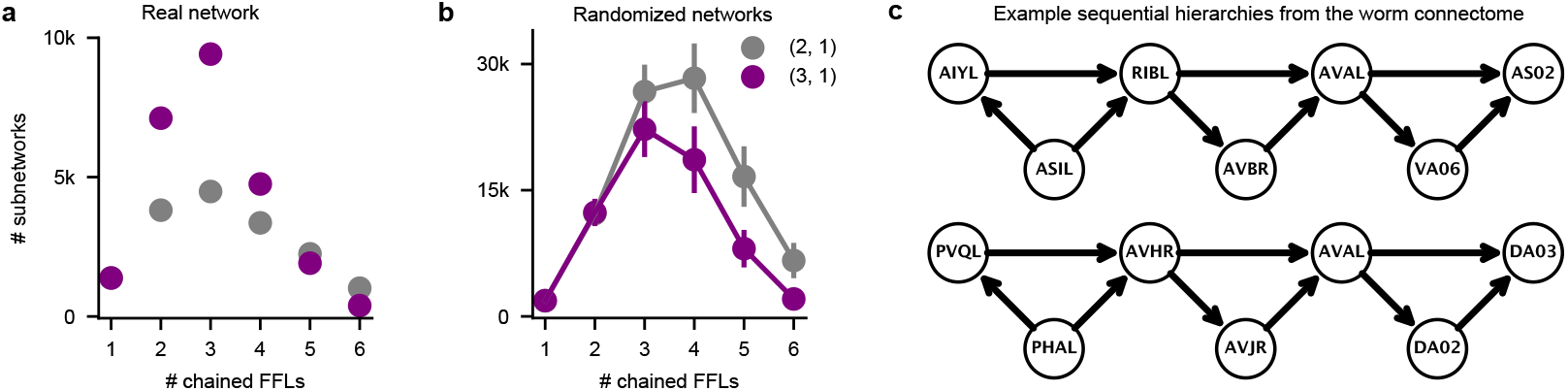
Sequential hierarchies of chain-length 3 are enriched in the *C. elegans* hermaphrodite connectome. **a**. Number of motifs matched by searching sequential hierarchies (purple) and intermediate node-chains (gray) of chain lengths 1-6 through the *C. elegans* hermaphrodite connectome [14] showing a preference of matching sequential hierarchies over intermediate node-chains from 2-4 chain lengths. **b**. Motif search of sequential hierarchies (purple) and intermediate node-chains (gray) of chain lengths 1-6 through randomized edge-swapped connectomes, showing a higher prevalence of intermediate node chains over sequential hierarchies in randomized networks longer than 2 chain-lengths. **c**. Two randomly chosen examples of length-3 sequential hierarchies matched in the *C. elegans* hermaphrodite connectome.

**Extended Data Fig. 3.**
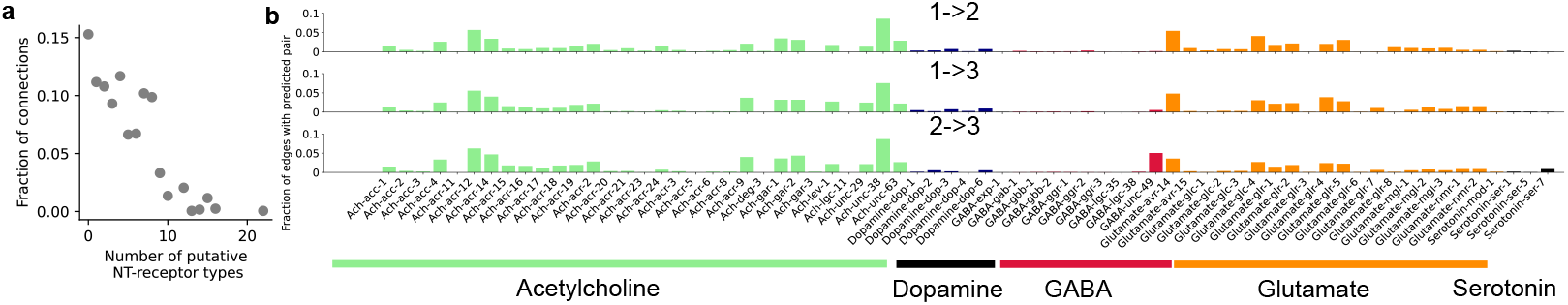
**a**. Histogram of the number of putative neurotransmitter (NT)-receptor pairs mapped to a given connection in the chemical connectome. **b**. Neurotransmitter ligand-receptor pairs for the feed-forward loop motif are more uniformly distributed with the exception of the GABA-unc-49, which is over-represented in the intermediate-node to output node connection.

**Extended Data Fig. 4.**
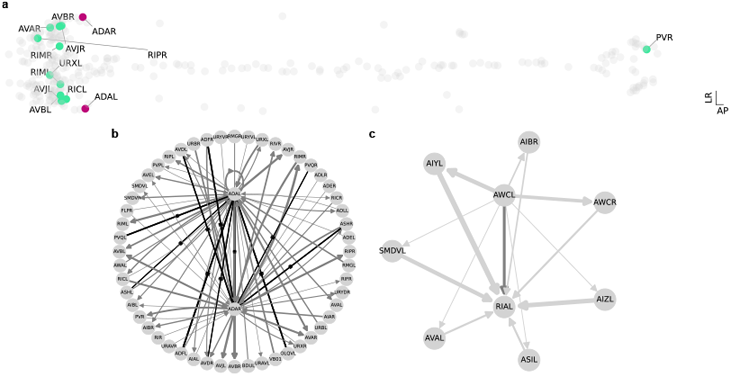
**a**. Neurons ADAL and ADAR (purple), and their connected neurons (light green) with >5 synapses in anatomical layout of all other neurons (gray) showing their relative positions in the nervous system. **b**. All direct connections to and from ADAL and ADAR shown as a shell plot. Gap junctions are shown in black with perpendicular marks on the edge. Chemical synapses in gray, with edge width representing number of synapses. **c**. Direct chemical synapses from AWCL to RIAL marked in dark gray, and indirect connections (of path-length 2) marked in light gray.

**Extended Data Fig. 5.**
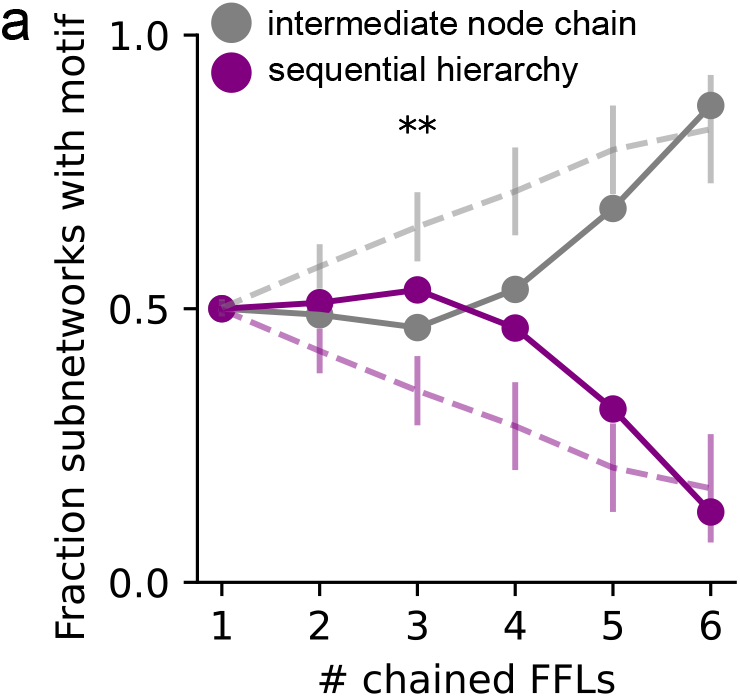
**a**. Motif search for sequential hierarchy and intermediate node-chains in the male *C. elegans* chemical connectome.

**Extended Data Fig. 6.**
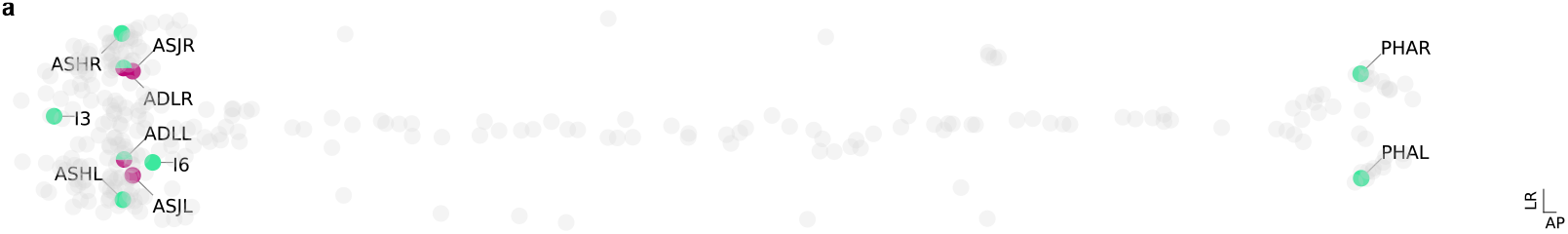
Integrating data from CENGEN into CeDNe. **a**. Anatomical layout showing the expression pattern of the sre-1, sre-2 and their intersection.

**Extended Data Fig. 7.**
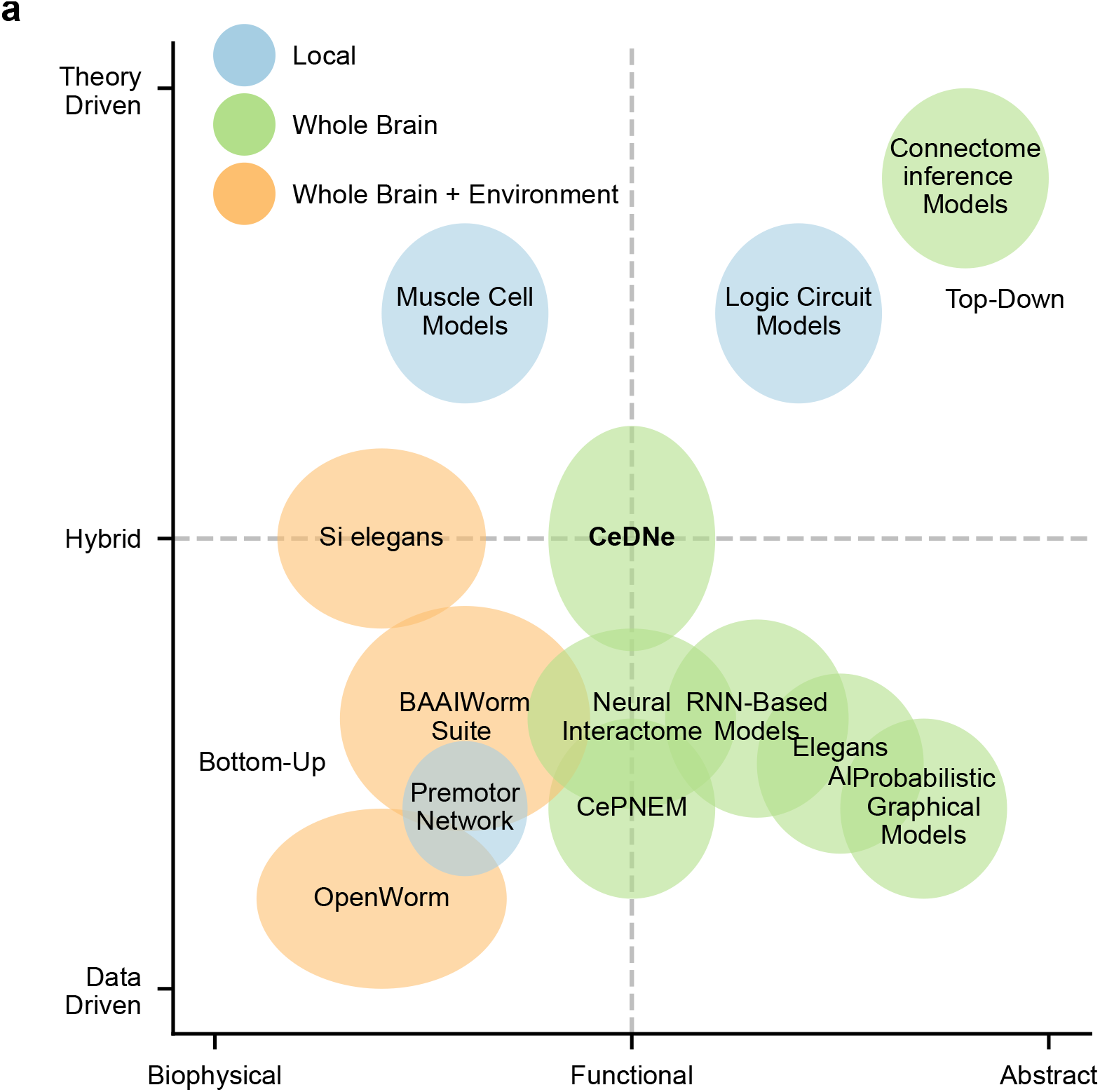
Position of CeDNe in the ecosystem of tools for *C. elegans* modeling frameworks. **a**. CeDNe approaches modeling the C. elegans nervous system from the top-down perspective and building functional, hybrid data-driven models.

## References

[1] Brenner, S.: NOBEL LECTURE: Nature’s Gift to Science. Bioscience Reports 23(5-6), 225–237 (2003) 10.1023/B:BIRE.0000019186.48208.f3. Accessed 2024-07-16

[2] Hammarlund, M., Hobert, O., Miller, D.M., Sestan, N.: The CeNGEN Project: The Complete Gene Expression Map of an Entire Nervous System. Neuron 99(3), 430–433 (2018) 10.1016/j.neuron.2018.07.042. Accessed 2023-07-05

[3] Taylor, S.R., Santpere, G., Weinreb, A., Barrett, A., Reilly, M.B., Xu, C., Varol, E., Oikonomou, P., Glenwinkel, L., McWhirter, R., Poff, A., Basavaraju, M., Rafi, I., Yemini, E., Cook, S.J., Abrams, A., Vidal, B., Cros, C., Tavazoie, S., Sestan, N., Hammarlund, M., Hobert, O., Miller, D.M.: Molecular topography of an entire nervous system. Cell 184(16), 4329–434723 (2021) 10.1016/j.cell.2021.06.023. Accessed 2023-07-05

[4] Altun, Zeynep F.: Neurotransmitter and Neuropeptide Receptors in Caenorhab ditis elegans 10.3908/WORMATLAS.5.202. Publisher: Wor mAtlas. Accessed 2025-11-02

[5] Wang, C., Vidal, B., Sural, S., Merritt, D.M., Toker, I.A., Vogt, M.C., Cros, C., Hobert, O.: A neurotransmitter atlas of the nervous system of C. elegans males and hermaphrodites (2024). 10.7554/eLife.95402.1. https://elifesciences.org/reviewed-preprints/95402v1 Accessed 2024-04-01

[6] Ripoll-Sánchez, L., Watteyne, J., Sun, H., Fernandez, R., Taylor, S.R., Weinreb, A., Bentley, B.L., Hammarlund, M., Miller, D.M., Hobert, O., Beets, I., Vértes, P.E., Schafer, W.R.: The neuropeptidergic connectome of C. elegans. Neuron 111(22), 3570–35895 (2023) 10.1016/j.neuron.2023.09.043. Accessed 2024-04-02

[7] Atanas, A.A., Kim, J., Wang, Z., Bueno, E., Becker, M., Kang, D., Park, J., Kramer, T.S., Wan, F.K., Baskoylu, S., Dag, U., Kalogeropoulou, E., Gomes, M.A., Estrem, C., Cohen, N., Mansinghka, V.K., Flavell, S.W.: Brain-wide representations of behavior spanning multiple timescales and states in C. elegans. Cell 186(19), 4134–415131 (2023) 10.1016/j.cell.2023.07.035. Accessed 2025-11-03

[8] Venkatachalam, V., Ji, N., Wang, X., Clark, C., Mitchell, J.K., Klein, M., Tabone, C.J., Florman, J., Ji, H., Greenwood, J., Chisholm, A.D., Srinivasan, J., Alkema, M., Zhen, M., Samuel, A.D.T.: Pan-neuronal imaging in roaming Caenorhabditis elegans. Proceedings of the National Academy of Sciences 113(8) (2016) 10.1073/pnas.1507109113. Accessed 2024-04-15

[9] Susoy, V., Hung, W., Witvliet, D., Whitener, J.E., Wu, M., Park, C.F., Graham, B.J., Zhen, M., Venkatachalam, V., Samuel, A.D.T.: Natural sensory context drives diverse brain-wide activity during C. elegans mating. Cell 184(20), 5122– 513717 (2021) 10.1016/j.cell.2021.08.024. Accessed 2024-04-15

[10] Kato, S., Kaplan, H., Schrödel, T., Skora, S., Lindsay, T., Yemini, E., Lockery, S., Zimmer, M.: Global Brain Dynamics Embed the Motor Command Sequence of Caenorhabditis elegans. Cell 163(3), 656–669 (2015) 10.1016/j.cell.2015.09.034. Accessed 2024-04-15

[11] Nguyen, J.P., Shipley, F.B., Linder, A.N., Plummer, G.S., Liu, M., Setru, S.U., Shaevitz, J.W., Leifer, A.M.: Whole-brain calcium imaging with cellular resolution in freely behaving Caenorhabditis elegans. Proceedings of the National Academy of Sciences 113(8) (2016) 10.1073/pnas.1507110112. Accessed 2024-04-15

[12] Liang, J., Moon, S., Moza, S., Lee, H.J., Eleftheriadis, P.E., Chen, J., Ge, M., Chen, M., Lu, H., Zhang, Y.: Aversive Learning Induces Context-Gated Global Reorganization of Neural Dynamics in Caenorhabditis elegans. Neuroscience (2025). 10.1101/2025.10.31.685731. http://biorxiv.org/lookup/doi/10.1101/2025.10.31.685731 Accessed 2025-11-02

[13] White, J.G., Southgate, E., Thomson, J.N., Brenner, S.: The structure of the nervous system of the nematode Caenorhabditis elegans. Philosophical Transactions of the Royal Society of London. Series B, Biological Sciences 314(1165), 1–340 (1986) 10.1098/rstb.1986.0056

[14] Cook, S.J., Jarrell, T.A., Brittin, C.A., Wang, Y., Bloniarz, A.E., Yakovlev, M.A., Nguyen, K.C.Q., Tang, L.T.-H., Bayer, E.A., Duerr, J.S., Bülow, H.E., Hobert, O., Hall, D.H., Emmons, S.W.: Whole-animal connectomes of both Caenorhabditis elegans sexes. Nature 571(7763), 63–71 (2019) 10.1038/s41586-019-1352-7. Accessed 2023-07-05

[15] Witvliet, D., Mulcahy, B., Mitchell, J.K., Meirovitch, Y., Berger, D.R., Wu, Y., Liu, Y., Koh, W.X., Parvathala, R., Holmyard, D., Schalek, R.L., Shavit, N., Chisholm, A.D., Lichtman, J.W., Samuel, A.D.T., Zhen, M.: Connectomes across development reveal principles of brain maturation. Nature 596(7871), 257–261 (2021) 10.1038/s41586-021-03778-8. Accessed 2024-04-03

[16] Haspel, G., Baker, B., Beets, I., Boyden, E.S., Brown, J., Church, G., Cohen, N., Colon-Ramos, D., Dyer, E., Fang-Yen, C., Flavell, S., Goodman, M.B., Hart, A.C., Izquierdo, E.J., Kagias, K., Lockery, S., Lu, Y., Marblestone, A., Matelsky, J., Mensh, B., Pereira, T.D., Pfister, H., Rajan, K., Rotstein, H.G., Scholz, M., Shaevitz, J.W., Shlizerman, E., Simeon, Q., Skuhersky, M.A., Tiruvadi, V., Venkatachalam, V., Wei, D., Wester, B., Yang, G.R., Yemini, E., Zimmer, M., Kording, K.P.: The time is ripe to reverse engineer an entire nervous system: simulating behavior from neural interactions. arXiv. Version Number: 5 (2023). 10.48550/ARXIV.2308.06578. https://arxiv.org/abs/2308.06578 Accessed 2024-11-19

[17] Alon, U.: Network motifs: theory and experimental approaches. Nature Reviews Genetics 8(6), 450–461 (2007) 10.1038/nrg2102. Accessed 2024-12-03

[18] Sarma, G.P., Lee, C.W., Portegys, T., Ghayoomie, V., Jacobs, T., Alicea, B., Cantarelli, M., Currie, M., Gerkin, R.C., Gingell, S., Gleeson, P., Gordon, R., Hasani, R.M., Idili, G., Khayrulin, S., Lung, D., Palyanov, A., Watts, M., Larson, S.D.: OpenWorm: overview and recent advances in integrative biological simulation of Caenorhabditis elegans. Philosophical Transactions of the Royal Society B: Biological Sciences 373(1758), 20170382 (2018) 10.1098/rstb.2017.0382. Accessed 2023-07-05

[19] Gleeson, P., Lung, D., Grosu, R., Hasani, R., Larson, S.D.: c302: a multiscale framework for modelling the nervous system of Caenorhabditis elegans. Philosophical Transactions of the Royal Society B: Biological Sciences 373(1758), 20170379 (2018) 10.1098/rstb.2017.0379. Accessed 2023-07-05

[20] Milo, R., Shen-Orr, S., Itzkovitz, S., Kashtan, N., Chklovskii, D., Alon, U.: Network Motifs: Simple Building Blocks of Complex Networks. Science 298(5594), 824–827 (2002) 10.1126/science.298.5594.824. Accessed 2025-10-21

[21] Bhatia, A., Moza, S., Bhalla, U.S.: Precise excitation-inhibition balance controls gain and timing in the hippocampus. eLife 8, 43415 (2019) 10.7554/eLife.43415. Accessed 2025-11-02

[22] Varshney, L.R., Chen, B.L., Paniagua, E., Hall, D.H., Chklovskii, D.B.: Structural Properties of the Caenorhabditis elegans Neuronal Network. PLoS Computational Biology 7(2), 1001066 (2011) 10.1371/journal.pcbi.1001066. Accessed 2024-12-04

[23] Fenyves, B.G., Szilágyi, G.S., Vassy, Z., Sőti, C., Csermely, P.: Synaptic polarity and sign-balance prediction using gene expression data in the Caenorhabditis elegans chemical synapse neuronal connectome network. PLOS Computational Biology 16(12), 1007974 (2020) 10.1371/journal.pcbi.1007974. Accessed 2025-05-24

[24] Gadhia, A., Barker, E., Morgan, A., Barclay, J.W.: Functional analysis of epilepsy-associated GABA_a_ receptor mutations using Caenorhabditis elegans. Epilepsia Open 9(4), 1458–1466 (2024) 10.1002/epi4.12982. Accessed 2025-10-21

[25] Dorkenwald, S., Matsliah, A., Sterling, A.R., Schlegel, P., Yu, S.-c., McKellar, C.E., Lin, A., Costa, M., Eichler, K., Yin, Y., Silversmith, W., Schneider-Mizell, C., Jordan, C.S., Brittain, D., Halageri, A., Kuehner, K., Ogedengbe, O., Morey, R., Gager, J., Kruk, K., Perlman, E., Yang, R., Deutsch, D., Bland, D., Sorek, M., Lu, R., Macrina, T., Lee, K., Bae, J.A., Mu, S., Nehoran, B., Mitchell, E., Popovych, S., Wu, J., Jia, Z., Castro, M.A., Kemnitz, N., Ih, D., Bates, A.S., Eckstein, N., Funke, J., Collman, F., Bock, D.D., Jefferis, G.S.X.E., Seung, H.S., Murthy, M., The FlyWire Consortium, Lenizo, Z., Burke, A.T., Willie, K.P., Serafetinidis, N., Hadjerol, N., Willie, R., Silverman, B., Ocho, J.A., Bañez, J., Candilada, R.A., Kristiansen, A., Panes, N., Yadav, A., Tancontian, R., Serona, S., Dolorosa, J.I., Vinson, K.J., Garner, D., Salem, R., Dagohoy, A., Skelton, J., Lopez, M., Capdevila, L.S., Badalamente, G., Stocks, T., Pandey, A., Akiatan, D.J., Hebditch, J., David, C., Sapkal, D., Monungolh, S.M., Sane, V., Pielago, M.L., Albero, M., Laude, J., Dos Santos, M., Vohra, Z., Wang, K., Gogo, A.M., Kind, E., Mandahay, A.J., Martinez, C., Asis, J.D., Nair, C., Patel, D., Manaytay, M., Tamimi, I.F.M., Lim, C.A., Ampo, P.L., Pantujan, M.D., Javier, A., Bautista, D., Rana, R., Seguido, J., Parmar, B., Saguimpa, J.C., Moore, M., Pleijzier, M.W., Larson, M., Hsu, J., Joshi, I., Kakadiya, D., Braun, A., Pilapil, C., Gkantia, M., Parmar, K., Vanderbeck, Q., Salgarella, I., Dunne, C., Munnelly, E., Kang, C.H., Lörsch, L., Lee, J., Kmecova, L., Sancer, G., Baker, C., Joroff, J., Calle, S., Patel, Y., Sato, O., Fang, S., Salocot, J., Salman, F., Molina-Obando, S., Brooks, P., Bui, M., Lichtenberger, M., Tamboboy, E., Molloy, K., Santana Cruz, A.E., Hernandez, A., Yu, S., Diwan, A., Patel, M., Aiken, T.R., Morejohn, S., Koskela, S., Yang, T., Lehmann, D., Chojetzki, J., Sisodiya, S., Koolman, S., Shiu, P.K., Cho, S., Bast, A., Reicher, B., Blanquart, M., Houghton, L., Choi, H., Ioannidou, M., Collie, M., Eckhardt, J., Gorko, B., Guo, L., Zheng, Z., Poh, A., Lin, M., Taisz, I., Murfin, W., Díez, S., Reinhard, N., Gibb, P., Patel, N., Kumar, S., Yun, M., Wang, M., Jones, D., Encarnacion-Rivera, L., Oswald, A., Jadia, A., Erginkaya, M., Drummond, N., Walter, L., Tastekin, I., Zhong, X., Mabuchi, Y., Figueroa Santiago, F.J., Verma, U., Byrne, N., Kunze, E., Crahan, T., Margossian, R., Kim, H., Georgiev, I., Szorenyi, F., Adachi, A., Bargeron, B., Stürner, T., Demarest, D., Gür, B., Becker, A.N., Turnbull, R., Morren, A., Sandoval, A., Moreno-Sanchez, A., Pacheco, D.A., Samara, E., Croke, H., Thomson, A., Laughland, C., Dutta, S.B., De Antón, P.G.A., Huang, B., Pujols, P., Haber, I., González-Segarra, A., Choe, D.T., Lukyanova, V., Mancini, N., Liu, Z., Okubo, T., Flynn, M.A., Vitelli, G., Laturney, M., Li, F., Cao, S., Manyari Diaz, C., Yim, H., Duc Le, A., Maier, K., Yu, S., Nam, Y., Bąba, D., Abusaif, A., Francis, A., Gayk, J., Huntress, S.S., Barajas, R., Kim, M., Cui, X., Sterne, G.R., Li, A., Park, K., Dempsey, G., Mathew, A., Kim, J., Kim, T., Wu, G.-t., Dhawan, S., Brotas, M., Zhang, C.-h., Bailey, S., Del Toro, A., Yang, R., Gerhard, S., Champion, A., Anderson, D.J., Behnia, R., Bidaye, S.S., Borst, A., Chiappe, E., Colodner, K.J., Dacks, A., Dickson, B., Garcia, D., Hampel, S., Hartenstein, V., Hassan, B., Helfrich-Forster, C., Huetteroth, W., Kim, J., Kim, S.S., Kim, Y.-J., Kwon, J.Y., Lee, W.-C., Linneweber, G.A., Maimon, G., Mann, R., Noselli, S., Pankratz, M., Prieto-Godino, L., Read, J., Reiser, M., Von Reyn, K., Ribeiro, C., Scott, K., Seeds, A.M., Selcho, M., Silies, M., Simpson, J., Waddell, S., Wernet, M.F., Wilson, R.I., Wolf, F.W., Yao, Z., Yapici, N., Zandawala, M.: Neuronal wiring diagram of an adult brain. Nature 634(8032), 124–138 (2024) 10.1038/s41586-024-07558-y. Accessed 2024-11-24

[26] Schlegel, P., Yin, Y., Bates, A.S., Dorkenwald, S., Eichler, K., Brooks, P., Han, D.S., Gkantia, M., Dos Santos, M., Munnelly, E.J., Badalamente, G., Ser ratosa Capdevila, L., Sane, V.A., Fragniere, A.M.C., Kiassat, L., Pleijzier, M.W., Stürner, T., Tamimi, I.F.M., Dunne, C.R., Salgarella, I., Javier, A., Fang, S., Perlman, E., Kazimiers, T., Jagannathan, S.R., Matsliah, A., Sterling, A.R., Yu, S.-c., McKellar, C.E., FlyWire Consortium, Kruk, K., Bland, D., Lenizo, Z., Burke, A.T., Willie, K.P., Bates, A.S., Serafetinidis, N., Hadjerol, N., Willie, R., Silverman, B., Ocho, J.A., Bañez, J., Candilada, R.A., Gager, J., Kristiansen, A., Panes, N., Yadav, A., Tancontian, R., Serona, S., Dolorosa, J.I., Vinson, K.J., Garner, D., Salem, R., Dagohoy, A., Skelton, J., Lopez, M., Stocks, T., Pandey, A., Akiatan, D.J., Hebditch, J., David, C., Sapkal, D., Monungolh, S.M., Sane, V., Pielago, M.L., Albero, M., Laude, J., Dos Santos, M., Deutsch, D., Vohra, Z., Wang, K., Gogo, A.M., Kind, E., Mandahay, A.J., Martinez, C., Asis, J.D., Nair, C., Patel, D., Manaytay, M., Lim, C.A., Ampo, P.L., Pantujan, M.D., Bautista, D., Rana, R., Seguido, J., Parmar, B., Saguimpa, J.C., Moore, M., Pleijzier, M.W., Larson, M., Hsu, J., Joshi, I., Kakadiya, D., Braun, A., Pilapil, C., Parmar, K., Vanderbeck, Q., Dunne, C., Munnelly, E., Kang, C.H., Lörsch, L., Lee, J., Kmecova, L., Sancer, G., Baker, C., Joroff, J., Calle, S., Patel, Y., Sato, O., Salocot, J., Salman, F., Molina-Obando, S., Bui, M., Lichtenberger, M., Tamboboy, E., Molloy, K., Santana-Cruz, A.E., Hernandez, A., Yu, S., Sorek, M., Diwan, A., Patel, M., Aiken, T.R., Morejohn, S., Koskela, S., Yang, T., Lehmann, D., Chojetzki, J., Sisodiya, S., Koolman, S., Shiu, P.K., Cho, S., Bast, A., Reicher, B., Blanquart, M., Houghton, L., Choi, H., Ioannidou, M., Collie, M., Eckhardt, J., Gorko, B., Guo, L., Zheng, Z., Poh, A., Lin, M., Taisz, I., Murfin, W., Díez, S., Reinhard, N., Gibb, P., Patel, N., Kumar, S., Yun, M., Wang, M., Jones, D., Encarnacion-Rivera, L., Oswald, A., Jadia, A., Erginkaya, M., Drummond, N., Walter, L., Tastekin, I., Zhong, X., Mabuchi, Y., Figueroa Santiago, F.J., Verma, U., Byrne, N., Kunze, E., Crahan, T., Margossian, R., Kim, H., Georgiev, I., Szorenyi, F., Adachi, A., Bargeron, B., Stürner, T., Demarest, D., Gür, B., Becker, A.N., Turnbull, R., Morren, A., Sandoval, A., Moreno-Sanchez, A., Pacheco, D.A., Samara, E., Croke, H., Thomson, A., Laughland, C., Dutta, S.B., De Antón, P.G.A., Huang, B., Pujols, P., Haber, I., González-Segarra, A., Lin, A., Choe, D.T., Lukyanova, V., Mancini, N., Liu, Z., Okubo, T., Flynn, M.A., Vitelli, G., Laturney, M., Li, F., Cao, S., Manyari-Diaz, C., Yim, H., Duc Le, A., Maier, K., Yu, S., Nam, Y., Bąba, D., Abusaif, A., Francis, A., Gayk, J., Huntress, S.S., Barajas, R., Kim, M., Cui, X., Sterling, A.R., Sterne, G.R., Li, A., Park, K., Dempsey, G., Mathew, A., Kim, J., Kim, T., Wu, G.-t., Dhawan, S., Brotas, M., Zhang, C.-h., Bailey, S., Del Toro, A., Lee, K., Macrina, T., Schneider-Mizell, C., Popovych, S., Ogedengbe, O., Yang, R., Halageri, A., Silversmith, W., Gerhard, S., Champion, A., Eckstein, N., Ih, D., Kemnitz, N., Castro, M., Jia, Z., Wu, J., Mitchell, E., Nehoran, B., Mu, S., Bae, J.A., Lu, R., Morey, R., Kuehner, K., Brittain, D., Jordan, C.S., Anderson, D.J., Behnia, R., Bidaye, S.S., Borst, A., Chiappe, E., Collman, F., Colodner, K.J., Dacks, A., Dickson, B., Funke, J., Garcia, D., Hampel, S., Hartenstein, V., Hassan, B., Helfrich-Forster, C., Huetteroth, W., Kim, J., Kim, S.S., Kim, Y.-J., Kwon, J.Y., Lee, W.-C., Linneweber, G.A., Maimon, G., Mann, R., Noselli, S., Pankratz, M., Prieto-Godino, L., Read, J., Reiser, M., Von Reyn, K., Ribeiro, C., Scott, K., Seeds, A.M., Selcho, M., Silies, M., Simpson, J., Waddell, S., Wernet, M.F., Wilson, R.I., Wolf, F.W., Yao, Z., Yapici, N., Zandawala, M., Costa, M., Seung, H.S., Murthy, M., Hartenstein, V., Bock, D.D., Jefferis, G.S.X.E.: Whole-brain annotation and multi-connectome cell typing of Drosophila. Nature 634(8032), 139–152 (2024) 10.1038/s41586-024-07686-5. Accessed 2024-11-24

[27] Jupyter, P., Bussonnier, M., Forde, J., Freeman, J., Granger, B., Head, T., Holdgraf, C., Kelley, K., Nalvarte, G., Osheroff, A., Pacer, M., Panda, Y., Perez, F., Ragan-Kelley, B., Willing, C.: Binder 2.0 - Reproducible, interactive, sharable environments for science at scale, Austin, Texas, pp. 113–120 (2018). 10.25080/Majora-4af1f417-011. https://doi.curvenote.com/10.25080/Majora4af1f417-011 Accessed 2025-11-02

[28] Kramer, T.S., Flavell, S.W.: Building and integrating brain-wide maps of nervous system function in invertebrates. Current Opinion in Neurobiology 86, 102868 (2024) 10.1016/j.conb.2024.102868. Accessed 2024-11-21

[29] Bazinet, V., Hansen, J.Y., Misic, B.: Towards a biologically annotated brain connectome. Nature Reviews Neuroscience 24(12), 747–760 (2023) 10.1038/s41583-023-00752-3. Accessed 2024-12-05

[30] Rübel, O., Tritt, A., Ly, R., Dichter, B.K., Ghosh, S., Niu, L., Baker, P., Soltesz, I., Ng, L., Svoboda, K., Frank, L., Bouchard, K.E.: The Neurodata Without Borders ecosystem for neurophysiological data science. eLife 11, 78362 (2022) 10.7554/eLife.78362. Accessed 2024-11-22

[31] Sprague, D.Y., Rusch, K., Dunn, R.L., Borchardt, J.M., Ban, S., Bubnis, G., Chiu, G.C., Wen, C., Suzuki, R., Chaudhary, S., Lee, H.J., Yu, Z., Dichter, B., Ly, R., Onami, S., Lu, H., Kimura, K.D., Yemini, E., Kato, S.: Unifying community-wide whole-brain imaging datasets enables robust automated neuron identification and reveals determinants of neuron positioning in C. elegans (2024). 10.1101/2024.04.28.591397. http://biorxiv.org/lookup/doi/10.1101/2024.04.28.591397 Accessed 2024-11-22

[32] Liu, H., Wu, T., Canales, X.G., Wu, M., Choi, M.-K., Duan, F., Calarco, J.A., Zhang, Y.: Forgetting generates a novel state that is reactivatable. Science Advances 8(6), 9071 (2022) 10.1126/sciadv.abi9071. Accessed 2025-03-09

[33] Szigeti, B., Gleeson, P., Vella, M., Khayrulin, S., Palyanov, A., Hokanson, J., Currie, M., Cantarelli, M., Idili, G., Larson, S.: OpenWorm: an open-science approach to modeling Caenorhabditis elegans. Frontiers in Computational Neuroscience 8 (2014) 10.3389/fncom.2014.00137. Accessed 2025-11-03

[34] Morrison, M., Young, L.-S.: A data-driven biophysical network model reproduces C. elegans premotor neural dynamics. arXiv. Version Number: 1 (2025). 10.48550/ARXIV.2501.00278. https://arxiv.org/abs/2501.00278 Accessed 2025-11-03

[35] Zhao, M., Wang, N., Jiang, X., Ma, X., Ma, H., He, G., Du, K., Ma, L., Huang, T.: An integrative data-driven model simulating C. elegans brain, body and environment interactions. Nature Computational Science 4(12), 978–990 (2024) 10.1038/s43588-024-00738-w. Accessed 2025-11-03

[36] Petrushin, A., Ferrara, L., Blau, A.: The Si elegans project at the interface of experimental and computational Caenorhabditis elegans neurobiology and behavior. Journal of Neural Engineering 13(6), 065001 (2016) 10.1088/1741-2560/13/6/065001. Accessed 2025-11-03

[37] Boyle, J.H., Cohen, N.: Caenorhabditis elegans body wall muscles are simple actuators. Biosystems 94(1-2), 170–181 (2008) 10.1016/j.biosystems.2008.05.025. Accessed 2025-11-03

[38] Mestre, G., Barbulescu, R., Oliveira, A.L., Silveira, L.M.: Modelling Neuronal Behaviour with Time Series Regression: Recurrent Neural Networks on C. Elegans Data. arXiv. 2107.06762 [q-bio] (2021). 10.48550/arXiv.2107.06762. http://arxiv.org/abs/2107.06762 Accessed 2025-11-03

[39] Bardozzo, F., Terlizzi, A., Simoncini, C., Lió, P., Tagliaferri, R.: Elegans-AI: How the connectome of a living organism could model artificial neural networks. Neurocomputing 584, 127598 (2024) 10.1016/j.neucom.2024.127598. Accessed 2025-11-03

[40] Liu, H., Kim, J., Shlizerman, E.: Functional connectomics from neural dynamics: probabilistic graphical models for neuronal network of Caenorhabditis elegans. Philosophical Transactions of the Royal Society B: Biological Sciences 373(1758), 20170377 (2018) 10.1098/rstb.2017.0377. Accessed 2025-11-03

[41] Kim, J., Leahy, W., Shlizerman, E.: Neural Interactome: Interactive Simulation of a Neuronal System. Frontiers in Computational Neuroscience 13, 8 (2019) 10.3389/fncom.2019.00008. Accessed 2025-11-03

[42] Yan, G., Vértes, P.E., Towlson, E.K., Chew, Y.L., Walker, D.S., Schafer, W.R., Barabási, A.-L.: Network control principles predict neuron function in the Caenorhabditis elegans connectome. Nature 550(7677), 519–523 (2017) 10.1038/nature24056. Accessed 2025-11-03

[43] Shukla, S., Dutta, S., Ganguly, U.: Design of Spiking Rate Coded Logic Gates for C. elegans Inspired Contour Tracking. In: Kůrková, V., Manolopoulos, Y., Hammer, B., Iliadis, L., Maglogiannis, I. (eds.) Artificial Neural Networks and Machine Learning – ICANN 2018 vol. 11139, pp. 273–283. Springer, Cham (2018). 10.1007/978-3-030-01418-6_27. Series Title: Lecture Notes in Computer Science. http://link.springer.com/10.1007/978-3-030-01418-627 Accessed2025 − 11 − 03

[44] Cordella, L.P., Foggia, P., Sansone, C., Vento, M.: A (sub)graph isomorphism algorithm for matching large graphs. IEEE Transactions on Pattern Analysis and Machine Intelligence 26(10), 1367–1372 (2004) 10.1109/TPAMI.2004.75. Accessed 2024-12-10

